# Solving the time-dependent protein distributions for autoregulated bursty gene expression using spectral decomposition

**DOI:** 10.1101/2023.11.21.568174

**Authors:** Bingjie Wu, James Holehouse, Ramon Grima, Chen Jia

## Abstract

In this study, we obtain an exact time-dependent solution of the chemical master equation (CME) of an extension of the two-state telegraph model describing bursty or non-bursty protein expression in the presence of positive or negative autoregulation. Using the method of spectral decomposition, we show that the eigenfunctions of the generating function solution of the CME are Heun functions, while the eigenvalues can be determined by solving a continued fraction equation. Our solution generalizes and corrects a previous time-dependent solution for the CME of a gene circuit describing non-bursty protein expression in the presence of negative autoregulation [“Exact time-dependent solutions for a self-regulating gene.” Phys. Rev. E 83: 062902 (2011)]. In particular, we clarify that the eigenvalues are generally not real as previously claimed. We also investigate the relationship between different types of dynamic behavior and the type of feedback, the protein burst size, and the gene switching rate.

## 1 Introduction

It is well known that often the expression of a gene is related to that of other genes. These gene-gene interactions are at the heart of gene regulatory networks [1]. Perhaps the simplest type of such interactions is autoregulation, whereby a protein influences its own transcription, leading to positive or negative feedback. This type of regulation is common, e.g. it is estimated that about 40% of *Escherichia coli*’s transcription factors self-regulate, mostly engaging in autorepression [2, 3].

Over the past 20 years, many stochastic models of autoregulation have been developed and their steady-state behavior has been studied using both simulations and theory. Within the continuous-time Markovian framework, some of these studies make headway by modelling gene state changes implicitly, and assuming protein numbers are real and positive (a continuum assumption) [4–8]. Other studies have tackled the more realistic problem where the change in molecule numbers of both gene and protein, when reactions occur, are integers [9–12]. Of the latter class of models, some assume that protein expression occurs one at a time [9, 10], whilst others assume that expression occurs in stochastic bursts [11, 12]. Another distinguishing feature is that some models assume that there is no change in the protein number when a protein copy binds to a gene or when it unbinds [9, 11], whilst others model the change explicitly [10, 12]. We note that the majority of these stochastic models have been solved exactly only in steady-state conditions. The exact time-dependent solution, being a much harder mathematical problem, has received considerably less attention — in [13], a model of non-bursty protein expression including negative autoregulation was purportedly solved exactly. A recent review of the various types of stochastic models of autoregulatory genetic circuits that also compares and contrasts their predictions can be found in [14].

In the present paper, we construct and exactly solve in time a stochastic model of bursty or non-bursty protein expression in the presence of positive or negative autoregulation, where both gene and protein numbers are modelled discretely. In Sec. 2, the reaction scheme and the chemical master equation (CME) describing the stochastic dynamics of the set of reactions are introduced. In Sec. 3, by means of the method of spectral decomposition, we show that the eigenfunctions of the time-dependent generating function solution of the CME are Heun functions, while the eigenvalues obey a continued fraction equation that is obtained by considering the holomorphism of the generating function. The accuracy of the solution is verified by stochastic simulations. Crucially, we also show that a previous time-dependent solution for the CME of a gene circuit describing non-bursty protein expression in the presence of negative autoregulation [13] is incorrect because the eigenvalues are generally not real as previously claimed. In Sec. 4, we investigate the relationship between five different types of dynamic behavior and the type of feedback, the protein burst size, and the gene switching rate. We conclude by a discussion in Sec. 5.

## 2 Model

Here we consider stochastic gene expression dynamics in a minimal coupled positive-plus-negative feedback loop with gene state switching, protein synthesis, and protein decay (Fig. 1). Let *G* and *G*^*\**^ denote the inactive and active states of the gene and let *P* denote the corresponding protein. In the active state *G*^*\**^, we assume that the protein is produced in a non-bursty (constitutive) or bursty manner. Both non-bursty and bursty gene expression are commonly observed in naturally occurring systems. Bursty protein synthesis is often due to rapid translation of protein from a single, short-lived mRNA molecule [15, 16]; if the mRNA lifetime is quite long (as common in mammalian cells [17]), then protein synthesis may appear non-bursty.

**Figure 1:**
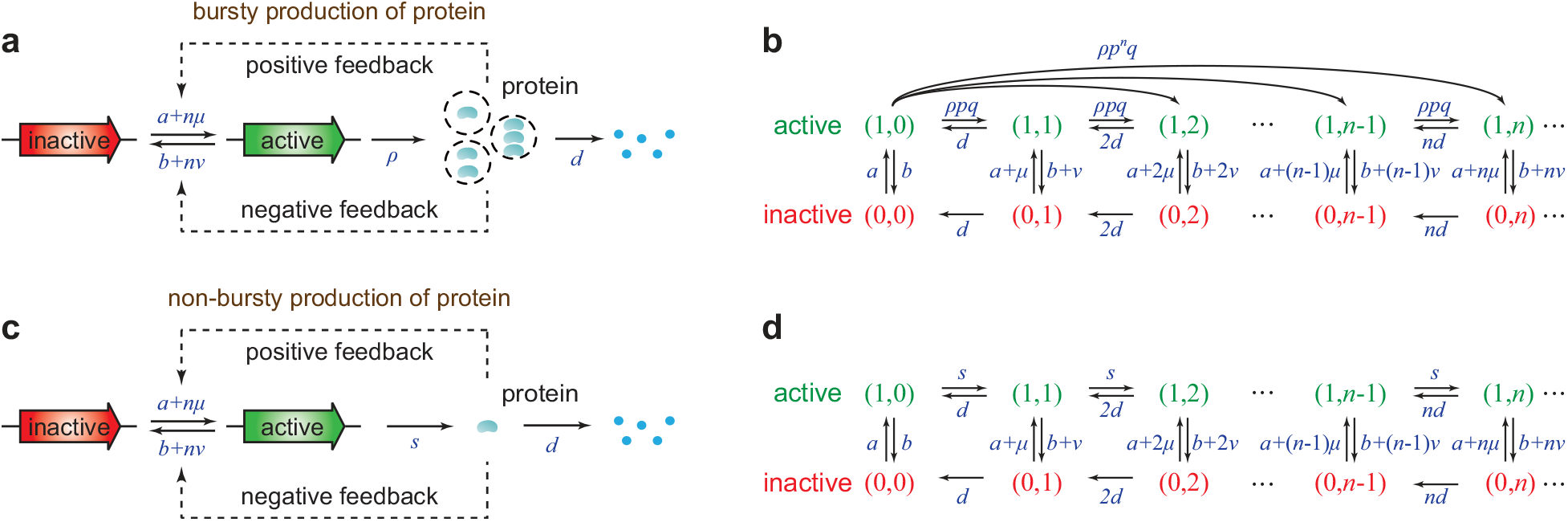
Models. **(a)** A minimal coupled gene circuit with positive-plus-negative feedback. When the gene is active, the protein is produced in a bursty manner with the burst size having a geometric distribution. **(b)** Transition diagram of the Markovian dynamics for the model illustrated in (a). Note that translational bursting can cause jumps from any microstate (1, *n*) to (1, *n* + *k*) (this is only shown for microstate (1, 0) in the figure but is also true for any other microstate (1, *n*)). **(c)** Same as (a) except that the protein is produced in a non-bursty manner when the gene is active, i.e. protein molecules are produced one at a time. **(d)** Transition diagram of the Markovian dynamics for the model illustrated in (c).

Let *n* be the copy number of the protein. In the bursty case, the reaction scheme underlying the coupled feedback loop is as follows (Fig. 1(a)):

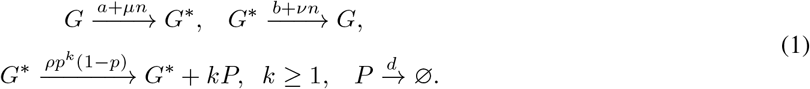

Due to feedback regulation, the protein number *n* will directly or indirectly affect the switching rates between the two gene states. Here *a* and *b* are the spontaneous switching rates, *μ* and *ν* characterize the strengths of positive and negative feedback loops, respectively, and *d* is the decay rate of the protein either due to protein degradation or due to dilution during cell division [18]. Specifically, the protein decay rate can be represented as *d* = log 2*/T*_*p*_ + log 2*/T*_*c*_, where *T*_*p*_ is the protein half-life and *T*_*c*_ is the cell cycle time. When the gene is in the active state *G*^*\**^, the synthesis of the protein occurs in bursts with frequency *ρ* and random size *k* sampled from a geometric distribution with parameter *p*, in agreement with experiments [19]. Note that the model considered here is more general than the classical model of autoregulatory feedback loops proposed by Kumar et al. [11]. In particular, the model describes a positive autoregulatory loop if the negative feedback strength *ν* vanishes, and it describes a negative autoregulatory loop if the positive feedback strength *μ* vanishes. In a coupled feedback loop, both *μ* and *ν* are nonzero. Coupled gene circuits widely exist in nature and they have been shown to be a crucial network motif to produce robust and tunable biological oscillations [20]. Biological examples of coupled feedback loops can be found in [20, 21].

The microstate of the gene can be represented by an ordered pair (*i, n*), where *i* is the gene state with *i* = 0, 1 corresponding to the inactive and active states, respectively, and *n* is the protein number. Let *p*_*i,n*_(*t*) denote the probability of having *n* protein molecules in an individual cell when the gene is in state *i*. Then the stochastic gene expression dynamics can be described by the Markov jump process shown in Fig. 1(b). The evolution of the Markovian dynamics is governed by the CMEs:

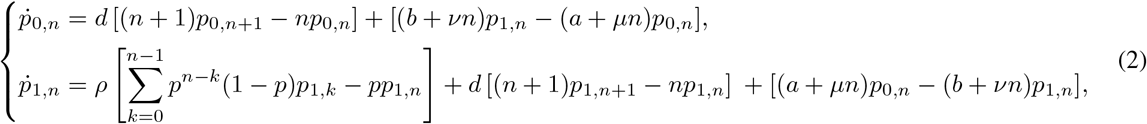

where the term involving *ρ* on the right-hand side represents protein synthesis, the terms involving *d* represent protein decay, and the terms involving *a, b, μ*, and *ν* represent gene state switching and feedback control.

In the non-bursty case, the reactions describing the coupled feedback loop are as follows (Fig. 1(c)):

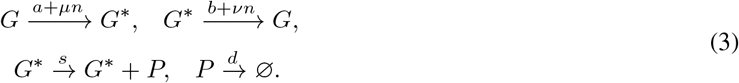

Here we assume that protein molecules are produced one at a time with rate *s* when the gene is active. The Markovian dynamics for this model is illustrated in Fig. 1(d). Note that the non-bursty model described by Eq. (3) is a limiting case of the bursty model described by Eq. (1) [22, 23]. Since the protein burst size is geometrically distributed, its expected value is given by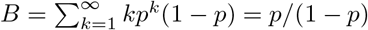. It is easy to see that when *ρ → ∞* and *B →* 0, while keeping *ρB* = *s* as constant, we have *p →* 0 and

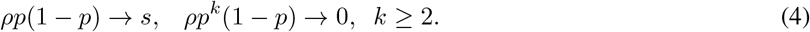

This shows that the bursty model reduces to the non-bursty one in the above limit. Hence in the following, we will first derive analytical results for the bursty model and then use them to obtain relevant results for the non-bursty model by taking the above limit.

We emphasize that the exact time-dependent distributions for the non-bursty model have been discussed in [13] in the special case of negative autoregulation (*μ* = *b* = 0). However, we find that solution given in [13] is both mathematically inconsistent and incomplete (the detailed reasons will be explained later). Here we generalize and correct the results obtained in previous studies by taking translational bursting and coupled feedback into account.

## 3 Solving the time-dependent master equation

### 3.1 Time-dependent solution for the bursty model

We first compute the time-dependent distributions of protein numbers for the bursty model. To this end, we define a pair of generating functions

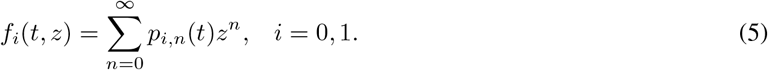

Moreover, let *p*_*n*_(*t*) = *p*_0,*n*_(*t*) + *p*_1,*n*_(*t*) denote the probability of having *n* protein molecules in an individual cell and let *f* (*t, z*) = *f*_0_(*t, z*) + *f*_1_(*t, z*) denote its generating function. Then Eq. (2) can be converted into the following set of partial differential equations (PDEs):

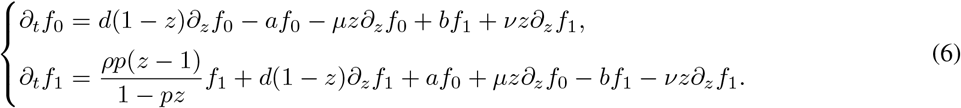

Following [13], we use the method of spectral decomposition to solve this set of PDEs. To this end, we assume that the generating functions *f*_*i*_ have the variable separation form of 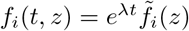. Here *λ* is called an *eigenvalue* of the PDEs given in Eq. (6) and 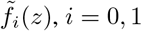 are called the corresponding *eigenfunctions*. Inserting it into Eq. (6) yields

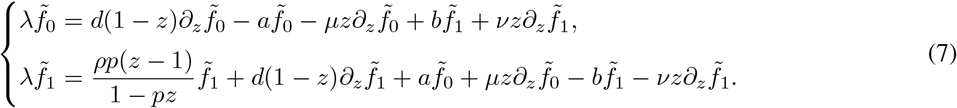

Moreover, let 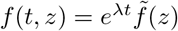 with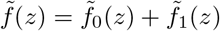. Adding the two identities in Eq. (7), we obtain

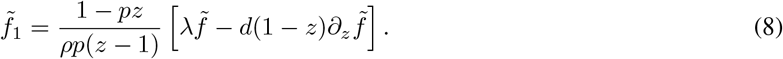

It then follows that

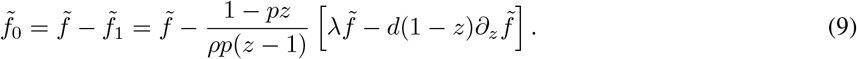

Substituting Eqs. (8) and (9) into the second row of Eq. (7) and eliminating 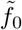 and 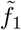, we find that 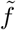 satisfies the following second-order ordinary differential equation (ODE):

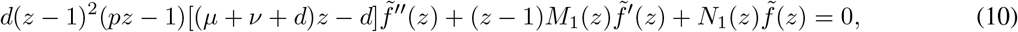

where

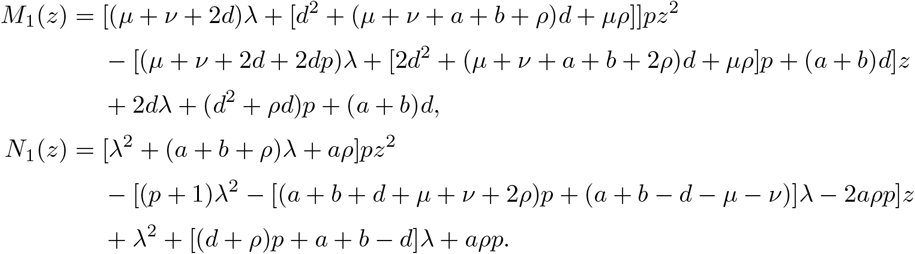

Note that the above ODE has four regular singularities 1, 1*/p, d/*(*μ* + *ν* + *d*), and *∞*. The crucial idea is to transform it into a Heun differential equation [13]. In fact, the regular singularities of the standard Heun differential equation are 0, 1, *ξ*, and *∞* for some constant *ξ* [24, Sec. 31.2]. Thus we need to make a variable transformation which maps *{d/*(*μ* + *ν* + *d*), 1, 1*/p, ∞}* onto *{*0, 1, *ξ, ∞}*. To this end, we introduce a new variable

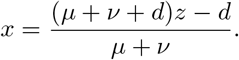

Moreover, let *h* be the function with the variable *x* associated with 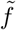, i.e.

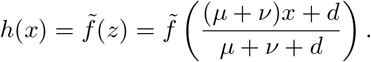

Then Eq. (10) can be transformed into the second-order ODE

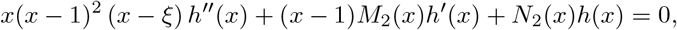

where *ξ* = (*μ* + *ν* + *d − dp*)*/*[(*μ* + *ν*)*p*] is a constant and

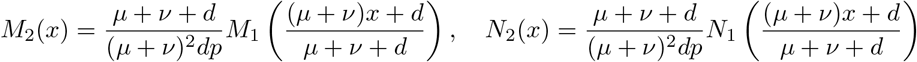

are functions of *x*. To proceed, we define a new function 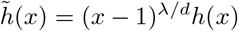. It is easy to check that 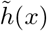 satisfies the following standard Heun differential equation:

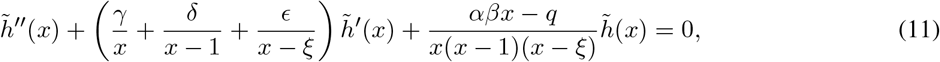

where

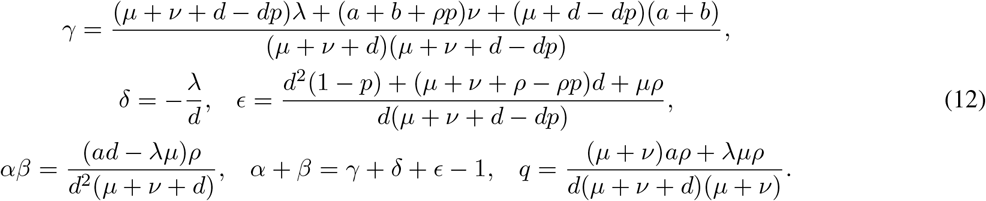

It is a classical result [25, 26] that the two general solutions of Eq. (11) are given by

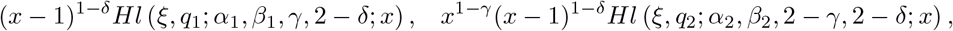

where *Hl* is the *local Heun function* (it is also referred to as the general Heun function [25]) and

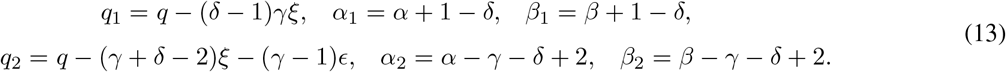

Hence the two linearly independent solutions of Eq. (10) are given by

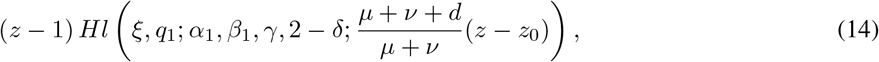

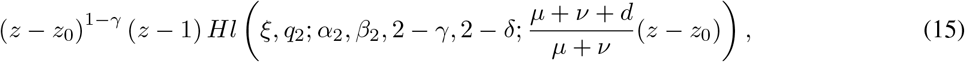

where *z*_0_ = *d/*(*μ* + *ν* + *d*).

We next make a crucial observation that in our model, the protein distribution *p*_*n*_(*t*) must decay exponentially with respect to the protein number *n* when *n ≥* 1 (see Supplementary Sec. 1 for the proof). In other words, *p*_*n*_(*t*) must have the following approximation:

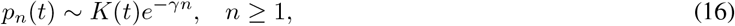

where *K*(*t*) is a constant depending on *t* and *γ >* 0 describes the decay rate of *p*_*n*_(*t*) with respect to *n*. This shows that

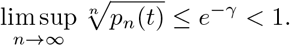

By the root test [27], the convergence radius of the power series given in Eq. (5) must be greater than 1. Since *z*_0_ *<* 1, the generating function *f* (*t, z*), as well as the function 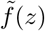, must be holomorphic at both *z* = *z*_0_ and *z* = 1. Recall that 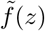 has two linearly independent solutions. In Supplementary Sec. 2, we prove that the solution given in Eq. (15) cannot be holomorphic at both *z* = *z*_0_ and *z* = 1. Hence we only need to consider the other solution given in Eq. (14).

Since the local Heun function *Hl*(*ξ, q*_1_; *α*_1_, *β*_1_, *γ*, 2 *− δ*; *x*) is holomorphic in the unit disk |*x*| *<* 1, it is clear that the solution given in Eq. (14) must be holomorphic at *z* = *z*_0_. On the other hand, the solution given in Eq. (14) is holomorphic at *z* = 1 if and only if the function *Hl*(*ξ, q*_1_; *α*_1_, *β*_1_, *γ*, 2 *− δ*; *x*) is holomorphic at *x* = 1. Note that the parameters *q*_1_, *α*_1_, *β*_1_, *γ*, and *δ* are all functions of the eigenvalue *λ* (see Eqs. (12) and (13)). It is an important property of the local Heun function [24, Sec. 31.4] that *Hl*(*ξ, q*_1_; *α*_1_, *β*_1_, *γ*, 2 *− δ*; *x*) is holomorphic at *x* = 1 if and only if *λ* satisfies the following continued fraction equation (also refer to [28]):

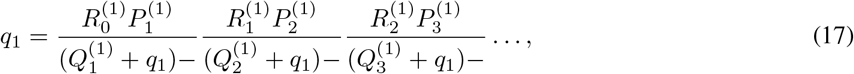

where

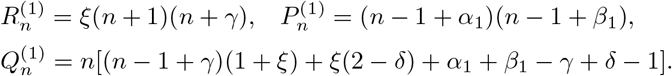

Here we have used the standard notation for continued fractions [24, Sec.1.12], i.e.

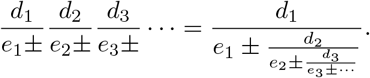

Solving Eq. (17) gives an infinite set of discrete values *λ*_*n*_, *n* = 1, 2, … and theses values are actually all nonzero eigenvalues of Eq. (6). Recall that the local Heun function *Hl*(*ξ, q*_1_; *α*_1_, *β*_1_, *γ*, 2 *− δ*; *x*) is called a *Heun function*, denoted by *Hf*, if it is holomorphic at *x* = 1 [24, Sec.31.4]. Then the eigenfunctions corresponding to all nonzero eigenvalues are given by

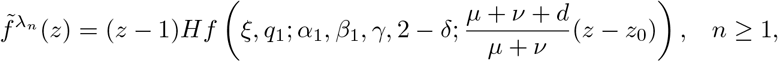

where *q*_1_, *α*_1_, *β*_1_, *γ*, and *δ* are all functions of the eigenvalue *λ*_*n*_.

We emphasize that solving the continued fraction equation can only give all nonzero eigenvalues. This is because for the zero eigenvalue *λ*_0_ = 0, it follows from Eq. (12) that *δ* = 0 and *αβx − q* = *aρ*(*x −* 1)*/*[*d*(*μ* + *ν* + *d*)]. In this case, the Heun differential equation given in Eq. (11) reduces to the hypergeometric differential equation

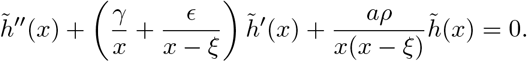

Similarly, by solving this equation, we find that the eigenfunction corresponding to the zero eigenvalue is given by

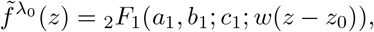

where

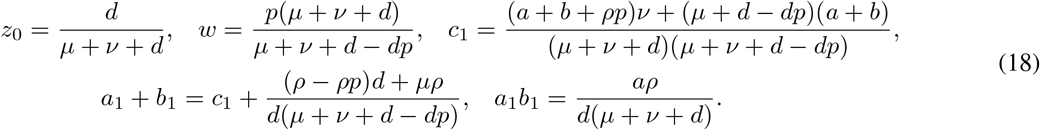

So far, we have derived the eigenfunctions corresponding to all eigenvalues. The eigenfunction corresponding to the zero eigenvalue is a hypergeometric function and the eigenfunctions corresponding to the nonzero eigenvalues have the form of Heun functions. By the Perron-Frobeniüs theorem, when the system is ergodic, one eigenvalue must be zero and the other eigenvalues must have negative real parts. Hence all eigenvalues can be arranged so that

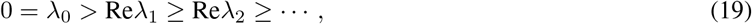

where Re(*z*) denotes the real part of *z*.

Recall that the stochastic dynamics of the coupled feedback loop is described by the Markov jump process illustrated in Fig. 1(b). In fact, all nonzero eigenvalues determined by solving Eq. (17) are exactly the same as the all nonzero eigenvalues of the generator matrix for the Markov jump process. Table 1 compares the first ten solutions of the continued fraction equation (truncated at *N* = 20 with *N* being the number of layers of the continued fraction equation) and the first ten nonzero eigenvalues of the generator matrix (truncated at *N* = 200 with *N* being the maximum protein number). Clearly, they coincides with each other perfectly and some eigenvalues may be complex numbers since the system is far from equilibrium.

**Table 1:** Comparison of the first ten nonzero eigenvalues computed using two different methods. All nonzero eigenvalues of the PDEs given in Eq. (6) are the solutions to the continued fraction equation given in Eq. (17). These eigenvalues are in excellent agreement with all nonzero eigenvalues of the generator matrix of the Markovian dynamics illustrated in Fig. 1(b). The parameters are chosen as *ρ* = 10, *B* = 1, *d* = 1, *μ* = 0.2, *ν* = 0.3, *a* = 1, *b* = 1.5.

| eigenvalues | $\lambda_1$ | $\lambda_2$ | $\lambda_3$ | $\lambda_4$ | $\lambda_5, \lambda_6$ | $\lambda_7, \lambda_8$ | $\lambda_9, \lambda_{10}$ |
| --- | --- | --- | --- | --- | --- | --- | --- |
| continued fraction method | -0.89 | -1.82 | -2.81 | -3.66 | $-4.56 \pm 0.33i$ | $-5.76 \pm 0.50i$ | $-6.99 \pm 0.57i$ |
| generator matrix method | -0.89 | -1.82 | -2.81 | -3.66 | $-4.56 \pm 0.33i$ | $-5.76 \pm 0.50i$ | $-7.00 \pm 0.55i$ |

Thus far, we have obtained all eigenvalues *λ*_*n*_, *n ≥* 0 and the corresponding eigenfunctions 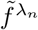 are given by

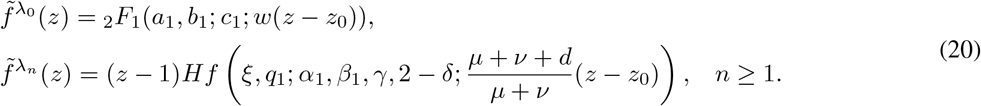

It follows from Eq. (8) that 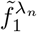 are given by

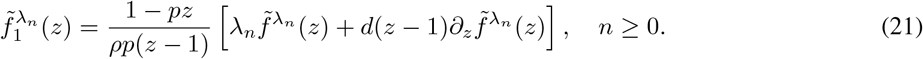

In this way, we can also determine 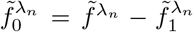. Finally, using the method of spectral decomposition, the generating functions *f*_*i*_ and *f* have the following series form:

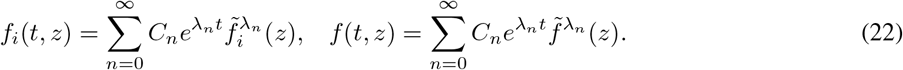

Then the time-dependent distribution of protein numbers can be recovered by taking the derivatives of the generating function *f* at *z* = 0, i.e.

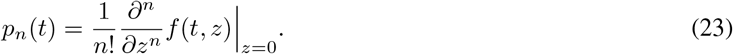

The remaining question is how to determine the coefficients *C*_*n*_ in Eq. (22) based on the initial conditions. To this end, it follows from Eq. (22) that for any complex numbers *z*_0_, *z*_1_, …, *z*_*M*_,

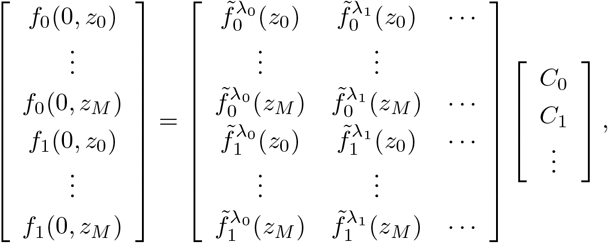

where 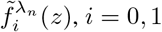 are determined by Eqs. (20) and (21), and 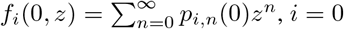, 1 are determined by the initial conditions. Note that this is an infinite dimensional system of linear equations with variables *C*_*n*_. In order to compute *C*_*n*_, we need to truncate the system at a large integer *M* . To do this, we use an approach similar to the inverse discrete Fourier transform. Let *M* be a positive integer and let *z*_*m*_ = *e*^2*πm***i***/M*^, 0 *≤ m ≤ M −* 1 be all *M* th roots of unity. Clearly, when *M ≫* 1, the coefficients *C*_*n*_ approximately satisfy the following set of linear equations:

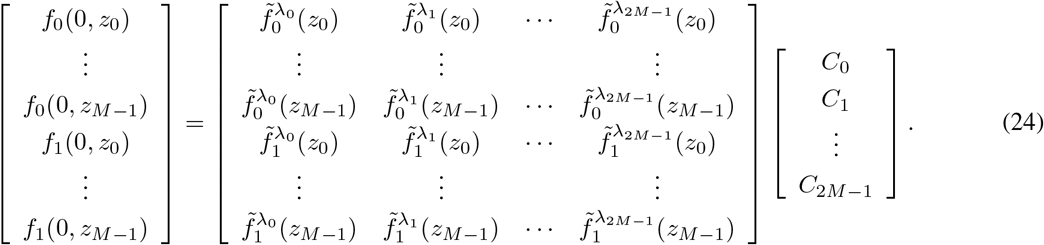

This is a finite dimensional system of linear equations with 2*M* variables, which can be solved either analytically or numerically (the analytical expression can be obtained using Cramer’s rule). However, the traditional numerical methods are not suitable for solving these equations since the coefficient matrix in Eq. (24) is too singular. Numerical computations indicate that traditional methods behave poorly when *M ≥* 20. To overcome this difficulty, note that the solution **x** to the linear equations *A***x** = **b** is also the solution to the optimization problem

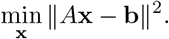

Following this idea, we transform Eq. (24) into the following equivalent optimization problem:

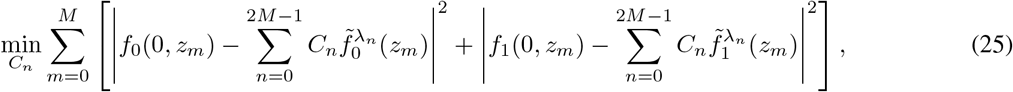

and then we use the MATLAB function “quadprog” to solve it. Thus far, we have obtained the full time-dependent solution for the bursty model. Note that when computing the eigenvalues *λ*_*n*_ and the coefficients *C*_*n*_, there are numerical steps involved and hence our solution is only semi-analytical. In the special case where the gene is unregulated (*μ* = *ν* = 0), both the eigenvalues and coefficients can be computed exactly, allowing us to obtain a full analytical expression of the time-dependent distribution (see Supplementary Secs. 3 and 4 for details).

To test our semi-analytical solution, we compare it with the numerical one obtained from the finite state projection algorithm (FSP) [29]. The results are shown in Fig. 2. When using FSP, we truncate the state space at a large protein number *N* and solve the truncated master equation numerically using the MATLAB function “ODE45”. The truncation size is chosen as *N* = 5*ρB/d*. Since *ρB/d* is the typical protein number in the active gene state, the probability that the protein number is outside this truncation size is very small and practically can always be ignored. As expected, the two solutions agree perfectly at all times. According to our simulations, for most sets of model parameters, the semi-analytical solution is computationally faster than FSP when the mean protein number is larger than *∼* 500; this is because the protein number needs to be truncated in FSP but does not in our solution.

**Figure 2:**
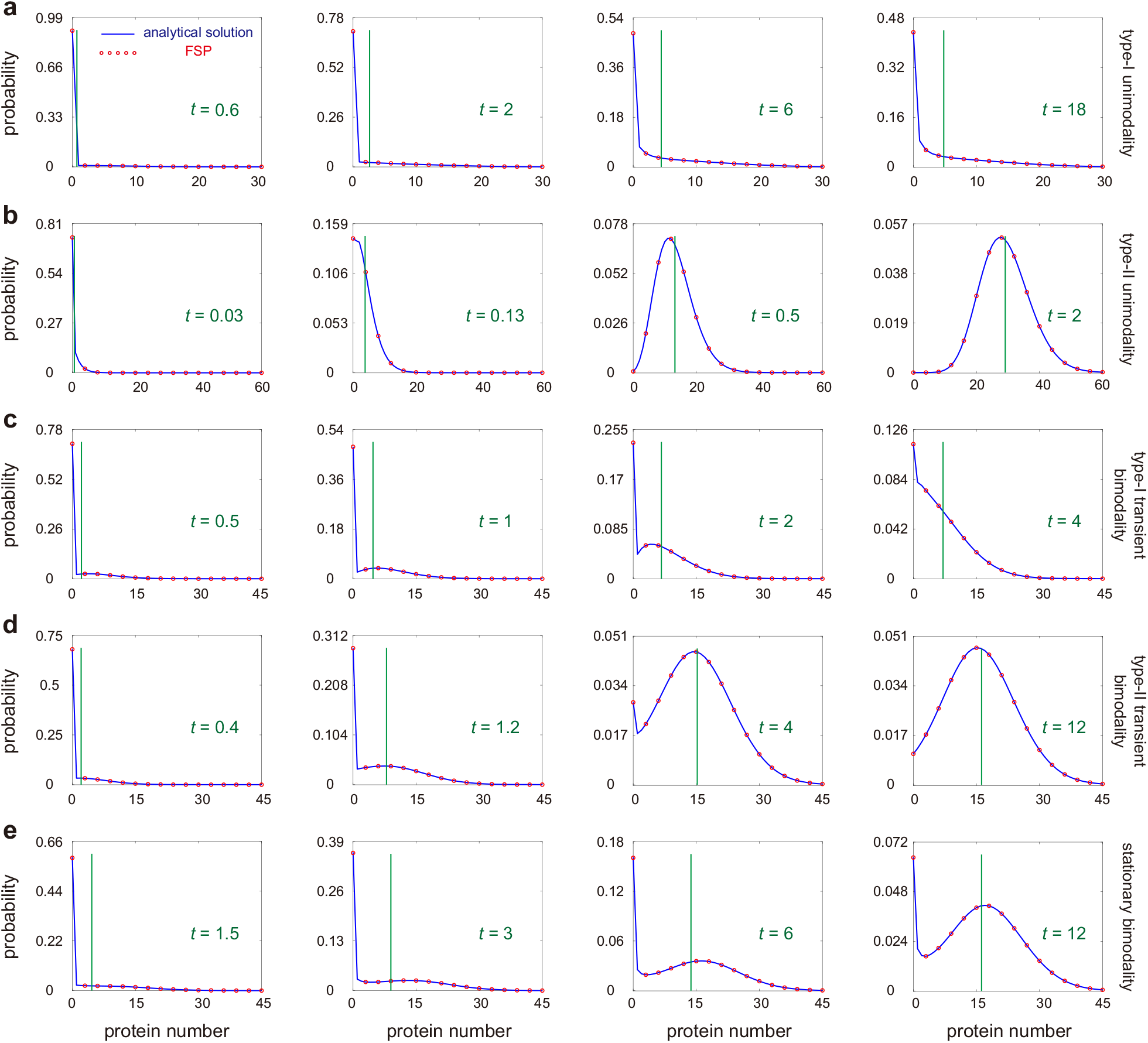
Comparison of the semi-analytical and numerical solutions for five different types of dynamic behavior at four time points. The blue curves show the semi-analytical distributions given in Eqs. (22) and (23), the red circles show the numerical ones obtained from FSP, and the green vertical lines show the mean protein numbers. In the semi-analytical solution, the eigenvalues *λ*_*n*_ are computed by solving Eq. (17) (the number of layers of the continued fraction equation is truncated at *N* = 200) and the coefficient *C*_*n*_ are computed by solving Eq. (25) for *M* = 50. Here we assume that initially the protein number is zero and the gene is off. **(a)**,**(b)** Unimodality. The distribution is unimodal at all times. **(a)** Type-I unimodality. The distribution peaks at zero at all times. The parameters are chosen as *ρ* = 35, *d* = 1, *μ* = 0.1, *ν* = 0.2, *a* = 0.2, *b* = 1, *B* = 1. **(b)** Type-II unimodality. The distribution has a zero peak at small times and has a nonzero peak at large times. The parameters are chosen as *ρ* = 35, *d* = 1, *μ* = 0.5, *ν* = 0.1, *a* = 80, *b* = 1, *B* = 1. **(c)**,**(d)** Transient bimodality. The distribution is unimodal at small and large times, and is bimodal at intermediate times. **(c)** Type-I transient bimodality. The distribution has a zero peak at large times. The parameters are chosen as *ρ* = 35, *d* = 1, *μ* = 0, *ν* = 0.3, *a* = 0.8, *b* = 0.1, *B* = 1. **(d)** Type-II transient bimodality. The distribution has a nonzero peak at large times. The parameters are chosen as *ρ* = 35, *d* = 1, *μ* = 0.2, *ν* = 0.2, *a* = 1.2, *b* = 1, *B* = 1. **(e)** Stationary bimodality. The distribution is unimodal at small times and is bimodal at large times. The parameters are chosen as *ρ* = 35, *d* = 1, *μ* = 0.3, *ν* = 0.2, *a* = 0.4, *b* = 1, *B* = 1.

### 3.2. Convergence to the steady-state solution

We next focus on the steady-state solution for the bursty model, which can be recovered from the time-dependent solution by taking *t → ∞*. We have seen that when the system is ergodic, one eigenvalue is *λ*_0_ = 0 and the other eigenvalues *λ*_*n*_, *n ≥* 1 have negative real parts. Hence the only term in Eq. (22) independent of time *t* is the first term and all other terms tend to zero exponentially fast as *t → ∞*. Then the steady-state generating function is given by

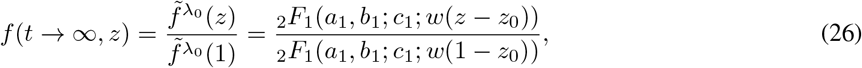

where we have used the fact that *f* (*t*, 1) = 1. Taking the derivatives of the generating function *f* at *z* = 0 yields the steady-state protein distribution

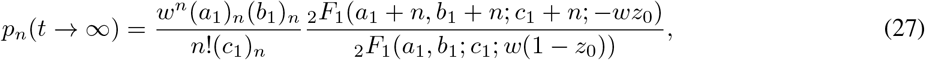

where (*a*)_*n*_ = *a*(*a* + 1) … (*a* + *n −* 1) denotes the Pochhammer symbol. Note that this is exactly the steady-state solution obtained previously in [22] which is a generalization of the special case of pure autoregulatory feedback studied earlier in [11].

### 3.3 Time-dependent solution for the non-bursty model

Next we focus on the time-dependent protein distributions for the non-bursty model. In this case, the generating functions *f*_*i*_ and *f* still have the form of

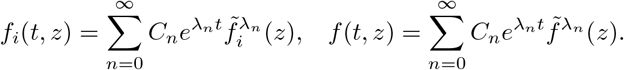

Recall that the non-bursty model given in Eq. (3) is a limiting case of the bursty model given in Eq. (1) when *ρ → ∞* and *B →* 0, while keeping *ρB* = *s* as constant. In this limit, for the zero eigenvalue *λ*_0_ = 0, the corresponding eigenfunction reduces to [24, Equation 13.18.2]

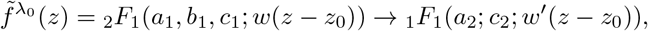

where _1_*F*_1_(*a*; *c*; *z*) is Kummer’s confluent hypergeometric function and

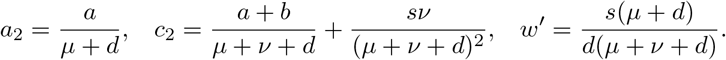

Moreover, in this limit, the Heun function *Hf* has the following limit (see Supplementary Sec. 5 for the proof):

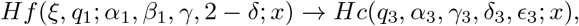

where *Hc* is the confluent Heun function [30] and

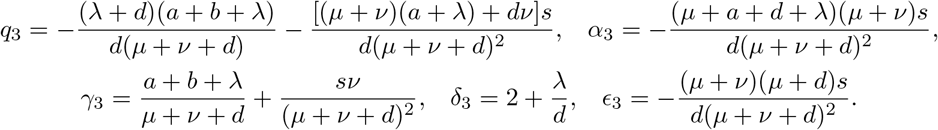

In addition, the continued fraction equation given in Eq. (17) reduces to

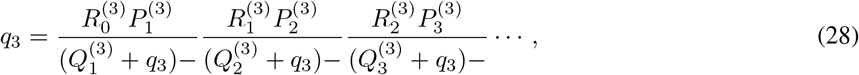

where

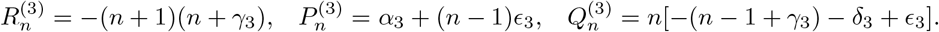

Similarly to the bursty case, all nonzero eigenvalues *λ*_*n*_, *n ≥* 1 can be obtained by solving Eq. (28), and the corresponding eigenfunctions reduce to

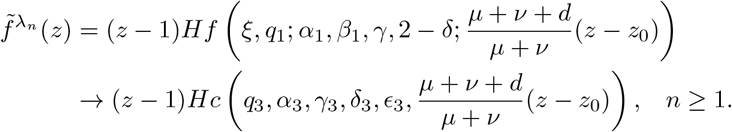

So far, we have obtained all the eigenvalues *λ*_*n*_ and eigenfunctions 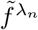. Similarly, by solving the optimization problem given in Eq. (25), we can compute all the coefficients *C*_*n*_. This gives the complete time-dependent solution for the non-bursty model.

We emphasize that for the non-bursty model, the time-dependent solution has been discussed in [13] in the special case of negative feedback loops (*μ* = *b* = 0). However, the solution derived in that paper is questionable due to the following two reasons. First, the authors did not show how to compute all the coefficients *C*_*n*_ based on the initial conditions. Second, the eigenvalues computed in [13] are incorrect. The authors claimed that the system has two families of eigenvalues (these eigenvalues were first reported in an earlier paper [31] by the same authors):

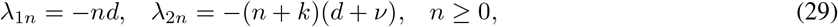

where *k* = *a/*(*d* + *ν*) + *sν/*(*d* + *ν*)^2^. However, the correct eigenvalues should be all the solutions to the continued fraction equation. Table 2 compares the correct eigenvalues obtained by solving Eq. (28) and the incorrect eigenvalues given in Eq. (29). We observe significant deviations between them; the former can be complex numbers, while the latter are always real. In Supplementary Sec. 6, we briefly explain how the false eigenvalues come from and further compare the true and false eigenvalues. Hence the solution derived in [13] is both incomplete and incorrect. In the present paper, we addressed the above two points.

**Table 2:** Comparison of the correct and incorrect eigenvalues for a negative autoregulatory loop. The first eight nonzero eigenvalues are computed using two different methods. The true eigenvalues are the solutions to Eq. (28), while the false eigenvalues are computed using Eq. (29). The parameters are chosen as *s* = 20, *d* = 1, *ν* = 0.1, *a* = 4, *μ* = *b* = 0.

| eigenvalues | $\lambda_1$ | $\lambda_2$ | $\lambda_3$ | $\lambda_4$ | $\lambda_5$ | $\lambda_6$ | $\lambda_7$ | $\lambda_8$ |
| --- | --- | --- | --- | --- | --- | --- | --- | --- |
| correct | -1.36 | -2.97 | -4.05+0.81i | -4.05-0.81i | -5.10+1.13i | -5.10-1.13i | -6.15+1.37i | -6.15-1.37i |
| incorrect | -1 | -2 | -3 | -4 | -5 | -5.82 | -6 | -6.92 |

## 4 Dynamical phase diagrams

We next investigate the shape of the time-dependent distribution for the bursty model. In what follows, we assume that the initial protein number is zero and the gene is initially in the inactive state [2]. This mimics the situation where the gene has been silenced by some repressor over a period of time such that all protein molecules have been removed via degradation. At time *t* = 0, the repressor is removed and we study how gene expression recovers. Under this initial condition, it is easy to see that *f*_0_(0, *z*) = 1 and *f*_1_(0, *z*) = 0.

In experiments, there are three commonly observed patterns for the protein distribution: a unimodal distribution with a zero peak (decaying distribution), a unimodal distribution with a nonzero peak (bell-shaped distribution), and a bimodal distribution with both a zero and a nonzero peak [32, 33]. Our model is capable of producing all these three distribution patterns (Fig. 2). Among the three patterns, the bimodal one attracts the most attention since it separates isogenic cells into two distinct phenotypes [34–36]. To further understand the shape of the time-dependent protein distribution, we classify the dynamic behavior of our model into five different phases: (i) the distribution is decaying at all times (Fig. 2(a)), (ii) the distribution is decaying at small times and is bell-shaped at large times (Fig. 2(b)), (iii) the distribution is decaying at small and large times, and is bimodal at intermediate times (Fig. 2(c)), (iv) the distribution is decaying at small times, is bimodal at intermediate times, and is bell-shaped at large times (Fig. 2(d)), and (v) the distribution is unimodal at small times and is bimodal at large times (Fig. 2(e)). To distinguish between them, we refer to (i) as type-I of unimodality (U1), to (ii) as type-II of unimodality (U2), to (iii) as type-I of transient bimodality (TB1), to (iv) as type-II of transient bimodality, and to (v) as stationary bimodality (SB). The semi-analytical solution and numerical solution obtained from FSP indicate that all the five dynamical phases can appear when model parameters are appropriately chosen (Fig. 2).

To determine the regions for the five phases in the parameter space, we illustrate the *a*-*b* phase diagrams of our model in Figs. 3 and 4. In all phase diagrams, there is a unique point separating the five phases; this is analogous to the triple point in physics and chemistry where all three phases (gas, liquid, and solid) of a substance coexist in thermodynamic equilibrium [37]. Intriguingly, we find that the regions for TB1 and TB2 are often not adjacent in the phase diagram and are separated by the SB region (see, e.g., the middle row of Fig. 4). This is why we distinguish type-I from type-II transient bimodality in the present paper. Fig. 3 shows the phase diagrams under different gene switching rates (slow switching, intermediate fast switching, and fast switching) and different feedback controls (positive feedback, coupled feedback, and negative feedback). We find that a positive feedback network fails to produce TB1, while a negative or coupled feedback network can produce all the five types of dynamical behavior. The region for transient and stationary bimodality (TB1, TB2, and SB) shrinks significantly as the switching rates increase. When the switching rates are relatively fast, the region for SB totally disappears in negative feedback networks, which agrees with the results found in [12, 38], and the region for TB1 also disappears in all three types of networks. This indicates that TB1 can only appear when the switching rates are relatively slow.

**Figure 3:**
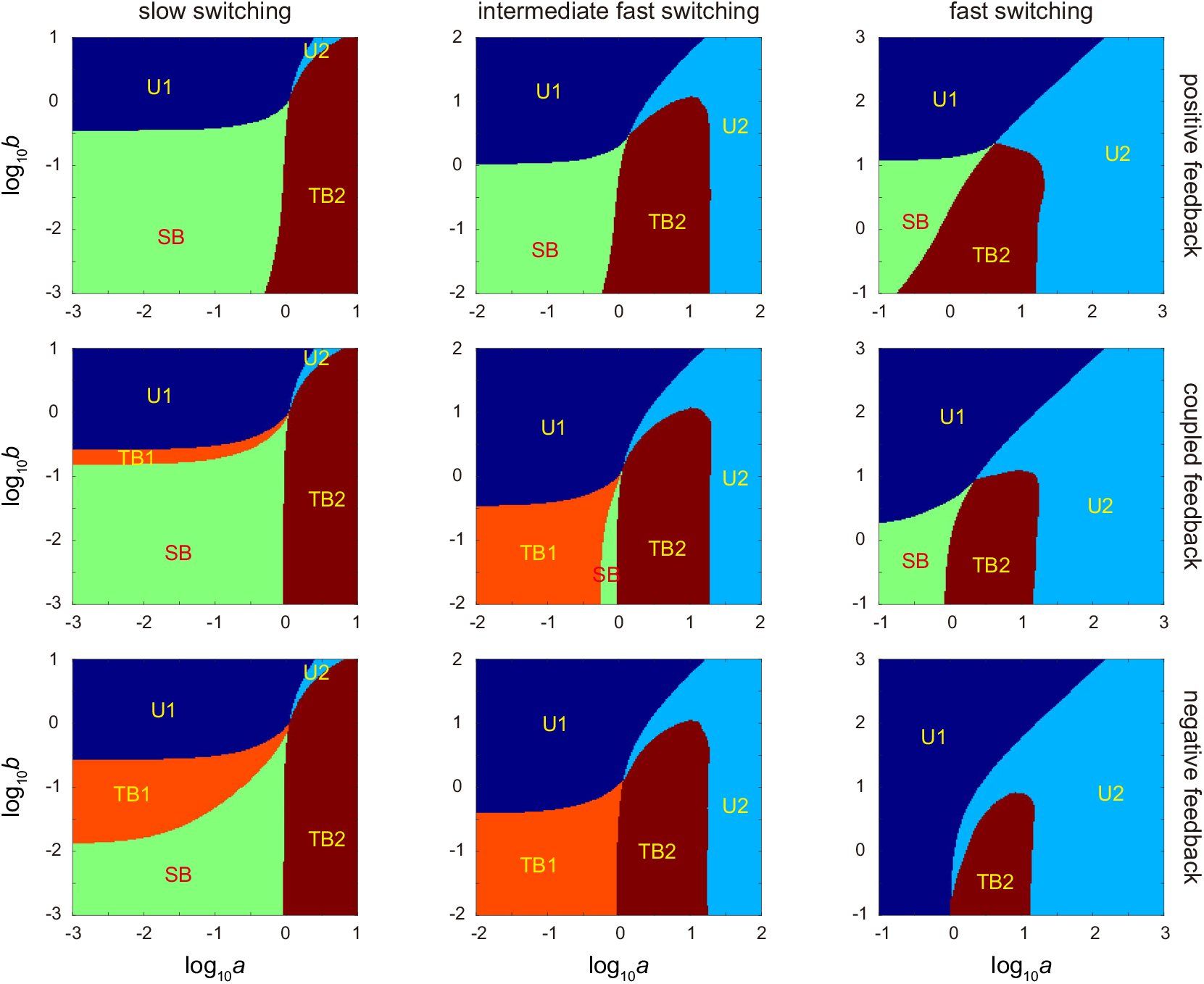
Dynamical phase diagrams in the *a*-*b* plane (here *a* and *b* are the spontaneous switching rates between the two gene states) under different switching rates and feedback controls. The system can produce five different types of dynamical behavior: type-I unimodality (U1, dark blue), type-II unimodality (U2, light blue), type-I transient bimodality (TB1, orange), type-II transient bimodality (TB2, brown), and stationary bimodality (SB, green). Note that in our model, the gene switching rates are *a* + *μn* and *b* + *νn*, with *μ* and *ν* being the strengths of positive and negative feedback loops, respectively. In slow switching conditions, the ranges of *a* and *b* are chosen to be 10^*−*3^ *−* 10^1^, and the feedback strengths are chosen as *μ* = 0.0225, *ν* = 0 for positive feedback, *μ* = 0.009, *ν* = 0.015 for coupled feedback, and *μ* = 0, *ν* = 0.0225 for negative feedback. In intermediate fast switching conditions, the ranges of *a* and *b* are chosen to be 10^*−*2^ *−* 10^2^, and the feedback strengths are chosen as *μ* = 0.225, *ν* = 0 for positive feedback, *μ* = 0.09, *ν* = 0.15 for coupled feedback, and *μ* = 0, *ν* = 0.225 for negative feedback. In fast switching conditions, the ranges of *a* and *b* are chosen to be 10^*−*1^ *−* 10^3^, and the feedback strengths are chosen as *μ* = 2.25, *ν* = 0 for positive feedback, *μ* = 0.9, *ν* = 1.5 for coupled feedback, and *μ* = 0, *ν* = 2.25 for negative feedback. The other parameters are chosen as *d* = 1, *B* = 1, *ρ* = 15.

**Figure 4:**
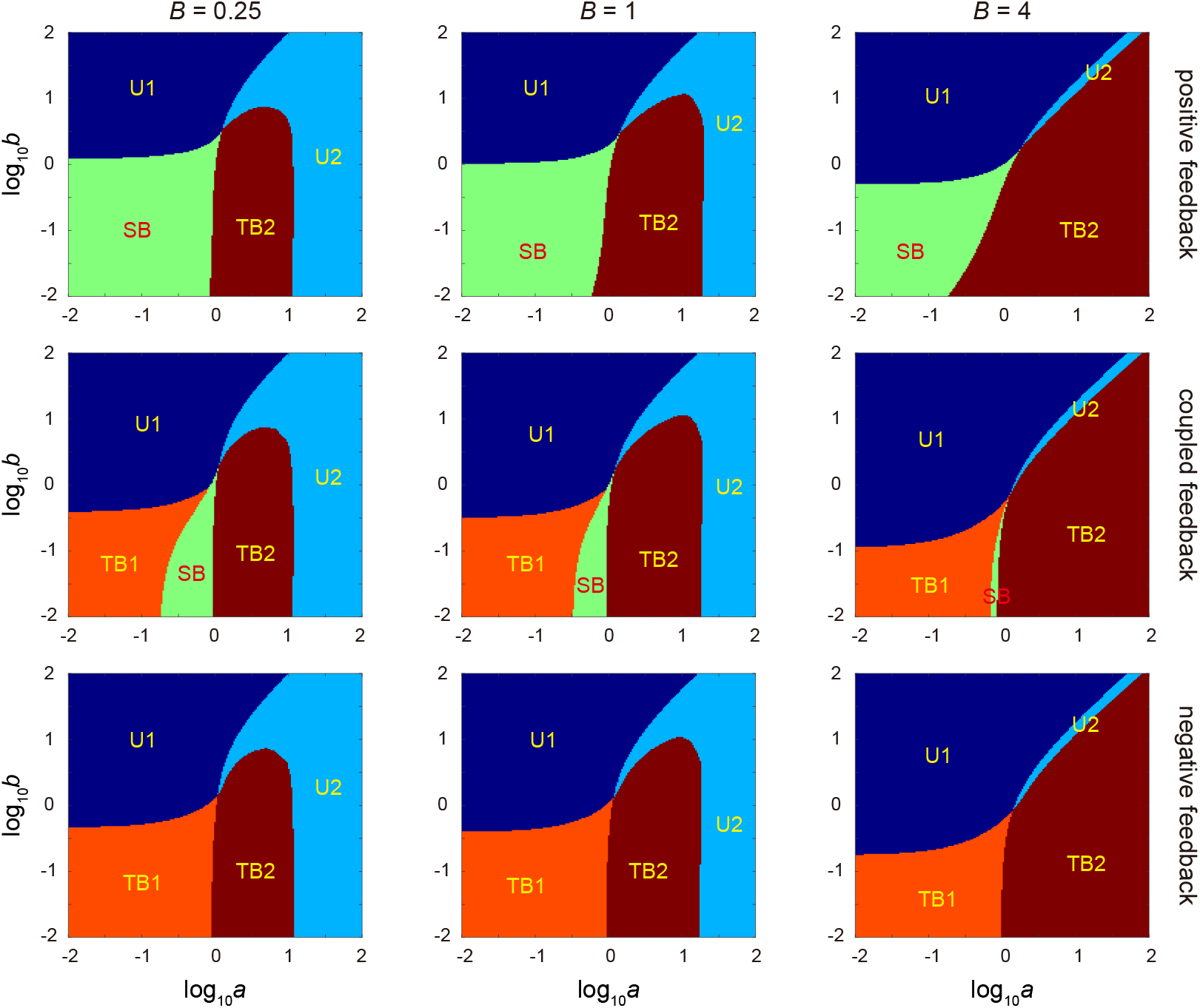
Dynamical phase diagrams in the *a*-*b* plane (here *a* and *b* are the spontaneous switching rates between the two gene states) under different burst sizes and feedback controls. The system can produce five different types of dynamical behavior: U1 (dark blue), U2 (light blue), TB1 (orange), TB2 (brown), and SB (green). The protein burst size is chosen as *B* = 0.25 (left), *B* = 1 (middle), and *B* = 4 (right). The feedback strengths are chosen as *μ* = 0.2, *ν* = 0 for positive feedback (upper), *μ* = *ν* = 0.1 for coupled feedback (middle), and *μ* = 0, *ν* = 0.2 for negative feedback (below). The other parameters are chosen as *d* = 1 and *ρ* is tuned so that the typical protein number *ρB/d* = 15 in the active gene state remains invariant.

We next focus on Fig. 4, which shows the phase diagrams under different protein burst sizes and feedback controls when the switching rates are not too small. Again, TB1 is not found in the positive feedback case and SB is not found in the negative feedback case (note that SB can be found when gene switching is very slow, as previously shown in Fig. 3). From the phase diagram, as the burst size *B* increases, the regions for TB1 and SB shrink, and the region for TB2 enlarges significantly in all three types of networks. In summary, we find that TB1 is mostly likely to occur in negative and coupled feedback networks when gene switching is not too fast; TB2 is mostly affected by the burst size; SB is sensitive to many factors and is mostly likely to occur in positive and coupled feedback networks when gene switching is relatively slow and the burst size is small.

## 5 Conclusion

In this work, we semi-analytically solved the CMEs and obtained the full time-dependence of a minimal coupled positive-plus-negative feedback loop with gene state switching, protein synthesis, and protein decay. This coupled gene circuit includes positive and negative autoregulatory feedback loops as special cases. In the active gene state, our model assumes that the protein is produced in a non-bursty or bursty manner. Following previous work [39], we transformed the CMEs satisfied by the protein distribution into the PDEs satisfied by the generating function. By using the method of spectral decomposition, we represented the protein distribution as a weighted sum of eigenvalue terms which only depends on time and eigenfunction terms which only depends on the spatial variable. We then make nontrivial spatial and functional transformations to obtain a Heun differential equation satisfied by the eigenfunctions.

Interestingly, the eigenfunctions are all local Heun functions, while the eigenvalues are those values such that these functions are holomorphic on the unit disk and can be determined by solving a continued fraction equation. In general, these eigenvalues cannot be computed in closed form. However, in the special case where the gene is unregulated, the eigenvalues can be solved exactly and the eigenfunctions reduce to Gaussian hypergeometric functions. We finally use a method similar to the inverse discrete Fourier transform to compute the weights of these eigenvalue and eigenfunction terms based on the initial conditions. In particular, our time-dependent solution generalizes the one obtained in [38] which can be applied only in fast switching conditions, and also generalizes and modifies the incomplete and incorrect solution obtained in [13] for the non-bursty case. The eigenvalues obtained in [13] are incorrect since they are all real, while the true eigenvalues can be complex numbers. The complex eigenvalues may be related to the oscillatory [40–42] and adaptation [43, 44] phenomena observed in negative feedback networks. Finally, we investigated the dynamical phase diagram of the coupled feedback loop by categorizing regions of parameter space according to five distinct types of time-evolution. This allowed us to deduce relationships between these phases and the type of feedback loop (positive, coupled, and negative), the burst size, and the speed of gene switching.

Our model has two notable limitations: (i) we assume that there is no change in the protein number during gene activation and inactivation. However, in reality, the protein number decreases by one when a protein copy binds to a gene and increases by one when unbinding occurs [10, 12]. In other words, the present model ignores the protein-gene binding fluctuations and hence it may not be accurate when the protein number is very small or when the feedback strength is very strong [12, 45]. The reason why we make this approximation is that if we ignore the binding fluctuations, then the differential equation satisfied by the eigenfunctions has four regular singularities and thus can be transformed into a Heun differential equation. However, if we take the binding fluctuations into account, then the differential equation will have five regular singularities and thus does not allow an existing special function representation; (ii) the model does not have an explicit cell cycle description. In reality, most proteins are not continuously degraded at some positive rate, but rather because their half-lives are often much longer than the cell cycle duration itself [17, 46, 47], they tend to accumulate during the cell cycle and then (approximately) half at cell division. Under some conditions, the modelling of protein degradation via a first-order reaction (as in our current model) well approximates the dilution due to cell division [48]. Note that our model with *d* = 0 can also be seen as predicting the time-dependent protein distribution within a cell cycle where *t* = 0 corresponds to the birth of a cell, but since there is no explicit modelling of cell division, it cannot predict how the distributions change from one generation to another. In the forthcoming Part II of this work, we will address some of these issues.

## Acknowledgements

The authors thank Dr. Youming Li for stimulating discussion. J. H. acknowledges support from National Science Foundation Grants EF-2133863 and DMR-191073. R. G. acknowledges support from the Leverhulme Trust (RPG-2020-327). C. J. acknowledges support from National Natural Science Foundation of China with grant No. U1930402 and No. 12271020.

## References

[1] Davidson, E. H. The regulatory genome: gene regulatory networks in development and evolution (Elsevier, 2010).

[2] Rosenfeld, N., Elowitz, M. B. & Alon, U. Negative autoregulation speeds the response times of transcription networks. Journal of molecular biology 323, 785–793 (2002).

[3] Shen-Orr, S. S., Milo, R., Mangan, S. & Alon, U. Network motifs in the transcriptional regulation network of Escherichia coli. Nature genetics 31, 64–68 (2002).

[4] Friedman, N., Cai, L. & Xie, X. S. Linking stochastic dynamics to population distribution: an analytical framework of gene expression. Phys. Rev. Lett. 97, 168302 (2006).

[5] Mackey, M. C., Tyran-Kaminska, M. & Yvinec, R. Dynamic behavior of stochastic gene expression models in the presence of bursting. SIAM J. Appl. Math. 73, 1830–1852 (2013).

[6] Bokes, P. & Singh, A. Protein copy number distributions for a self-regulating gene in the presence of decoy binding sites. PloS one 10, e0120555 (2015).

[7] Pájaro, M., Alonso, A. A. & Vázquez, C. Shaping protein distributions in stochastic self-regulated gene expression networks. Physical Review E 92, 032712 (2015).

[8] Jia, C., Zhang, M. Q. & Hong, Q. Emergent Lévy behavior in single-cell stochastic gene expression. Phys. Rev. E 96, 040402(R) (2017).

[9] Hornos, J. E. et al. Self-regulating gene: an exact solution. Physical Review E 72, 051907 (2005).

[10] Grima, R., Schmidt, D. R. & Newman, T. J. Steady-state fluctuations of a genetic feedback loop: An exact solution. The Journal of chemical physics 137 (2012).

[11] Kumar, N., Platini, T. & Kulkarni, R. V. Exact distributions for stochastic gene expression models with bursting and feedback. Phys. Rev. Lett. 113, 268105 (2014).

[12] Jia, C. & Grima, R. Small protein number effects in stochastic models of autoregulated bursty gene expression. J. Chem. Phys. 152 (2020).

[13] Ramos, A. F., Innocentini, G. C. & Hornos, J. E. M. Exact time-dependent solutions for a self-regulating gene. Phys. Rev. E 83, 062902 (2011).

[14] Holehouse, J., Cao, Z. & Grima, R. Stochastic modeling of autoregulatory genetic feedback loops: a review and comparative study. Biophysical Journal 118, 1517–1525 (2020).

[15] Paulsson, J. Models of stochastic gene expression. Phys. Life Rev. 2, 157–175 (2005).

[16] Jia, C. Simplification of Markov chains with infinite state space and the mathematical theory of random gene expression bursts. Phys. Rev. E 96, 032402 (2017).

[17] Schwanhäusser, B. et al. Global quantification of mammalian gene expression control. Nature 473, 337 (2011).

[18] Paulsson, J. & Ehrenberg, M. Noise in a minimal regulatory network: plasmid copy number control. Q. Rev. Biophys. 34, 1–59 (2001).

[19] Cai, L., Friedman, N. & Xie, X. S. Stochastic protein expression in individual cells at the single molecule level. Nature 440, 358–362 (2006).

[20] Tsai, T. Y.-C. et al. Robust, tunable biological oscillations from interlinked positive and negative feedback loops. Science 321, 126–129 (2008).

[21] Liu, P., Yuan, Z., Wang, H. & Zhou, T. Decomposition and tunability of expression noise in the presence of coupled feedbacks. Chaos 26, 043108 (2016).

[22] Jia, C., Yin, G. G., Zhang, M. Q. et al. Single-cell stochastic gene expression kinetics with coupled positive-plus-negative feedback. Phys. Rev. E 100, 052406 (2019).

[23] Jia, C. & Li, Y. Analytical time-dependent distributions for gene expression models with complex promoter switching mechanisms. SIAM J. Appl. Math. 83, 1572–1602 (2023).

[24] Olver, F. W. J. et al. NIST Digital Library of Mathematical Functions. http://dlmf.nist.gov/, Release 1.0.17 of 2017–12-22 (2017).

[25] Ronveaux, A. Heun’s differential equations (The Clarendon Press Oxford University Press, New York, 1995).

[26] Maier, R. The 192 solutions of the Heun equation. Mathematics of Computation 76, 811–843 (2007).

[27] Rudin, W. Principles of mathematical analysis (McGraw Hill, 1953).

[28] Holehouse, J. & Moran, J. Exact time-dependent dynamics of discrete binary choice models. Journal of Physics: Complexity 3, 035005 (2022).

[29] Munsky, B. & Khammash, M. The finite state projection algorithm for the solution of the chemical master equation. J. Chem. Phys. 124, 044104 (2006).

[30] Motygin, O. V. On evaluation of the confluent Heun functions. In 2018 Days on Diffraction (DD), 223–229 (IEEE, 2018).

[31] Ramos, A. F. & Hornos, J. E. Symmetry and stochastic gene regulation. Phys. Rev. Lett. 99, 108103 (2007).

[32] Munsky, B., Neuert, G. & Van Oudenaarden, A. Using gene expression noise to understand gene regulation. Science 336, 183–187 (2012).

[33] Jiao, F., Sun, Q., Tang, M., Yu, J. & Zheng, B. Distribution modes and their corresponding parameter regions in stochastic gene transcription. SIAM J. Appl. Math. 75, 2396–2420 (2015).

[34] Kærn, M., Elston, T. C., Blake, W. J. & Collins, J. J. Stochasticity in gene expression: from theories to phenotypes. Nat. Rev. Genet. 6, 451–464 (2005).

[35] Jia, C., Qian, M., Kang, Y. & Jiang, D. Modeling stochastic phenotype switching and bet-hedging in bacteria: stochastic nonlinear dynamics and critical state identification. Quant. Biol. 2, 110–125 (2014).

[36] Thomas, P., Popovic, N. & Grima, R. Phenotypic switching in gene regulatory networks. Proc. Natl. Acad. Sci. USA 111, 6994–6999 (2014).

[37] McNaught, A. D., Wilkinson, A. et al. Compendium of chemical terminology, vol. 1669 (Blackwell Science Oxford, 1997).

[38] Jia, C. & Grima, R. Dynamical phase diagram of an auto-regulating gene in fast switching conditions. J. Chem. Phys. 152 (2020).

[39] Berg, O. G. A model for the statistical fluctuations of protein numbers in a microbial population. J. Theor. Biol. 71, 587–603 (1978).

[40] Novák, B. & Tyson, J. J. Design principles of biochemical oscillators. Nat. Rev. Mol. Cell Biol. 9, 981–991 (2008).

[41] Jia, C., Zhang, M. Q. & Qian, H. Analytic theory of stochastic oscillations in single-cell gene expression. arXiv preprint arXiv:1909.09769 (2019).

[42] Gupta, A. & Khammash, M. Frequency spectra and the color of cellular noise. Nat. Commun. 13, 4305 (2022).

[43] Ma, W., Trusina, A., El-Samad, H., Lim, W. A. & Tang, C. Defining network topologies that can achieve biochemical adaptation. Cell 138, 760–773 (2009).

[44] Jia, C. & Qian, M. Nonequilibrium enhances adaptation efficiency of stochastic biochemical systems. PloS one 11, e0155838 (2016).

[45] Holehouse, J. & Grima, R. Revisiting the reduction of stochastic models of genetic feedback loops with fast promoter switching. Biophys. J. 117, 1311–1330 (2019).

[46] Christiano, R., Nagaraj, N., Fröhlich, F. & Walther, T. C. Global proteome turnover analyses of the yeasts S. cerevisiae and S. pombe. Cell Rep. 9, 1959–1965 (2014).

[47] Jia, C. & Grima, R. Frequency domain analysis of fluctuations of mRNA and protein copy numbers within a cell lineage: theory and experimental validation. Phys. Rev. X 11, 021032 (2021).

[48] Beentjes, C. H., Perez-Carrasco, R. & Grima, R. Exact solution of stochastic gene expression models with bursting, cell cycle and replication dynamics. Phys. Rev. E 101, 032403 (2020).

