## Supplementary Material for "Solving the time-dependent protein distributions for autoregulated bursty gene expression using spectral decomposition"

#### Contents

**1 Proof of Eq. (16) in the main text** **2**

**2 Proof that Eq. (15) in the main text cannot be holomorphic at both $z = z_0$ and $z = 1$** **2**

**3 Analytical time-dependent solution for unregulated genes** **3**

**4 Proof of the results in Section 3** **4**

**5 Reduction of the Heun function to the confluent Heun function** **8**

**6 Origin of the incorrect eigenvalues and comparison with FSP** **9**

### 1 Proof of Eq. (16) in the main text

Here we will prove that the protein distribution $p_n(t)$ decays exponentially with respect to the protein number $n$ when $n \gg 1$. To this end, we first consider the simple case where the gene is always active. In this case, the reaction scheme describing the gene expression dynamics is as follows:

$$G^* \xrightarrow{\rho p^k(1-p)} G^* + kP, \quad k \geq 1, \quad P \xrightarrow{d} \varnothing. \tag{1}$$

Without loss generality, we assume that the initial protein number is zero. Recall that the exact time-dependent generating function $f$ of the above model has been derived in [1, 2] and is given by

$$f(t,z) = \left[\frac{1 - B(z-1)e^{-dt}}{1 + B - Bz}\right]^{\rho/d}.$$

It is easy to check that the convergence radius of $f$ at $z = 0$ is $(1+B)/B = 1/p$. By the root test of power series, the distribution $\tilde{p}_n(t)$ of the above model must satisfy

$$\limsup_{n\to\infty} \sqrt[n]{\tilde{p}_n(t)} = p < 1.$$

This implies that there exists $C(t) \geq 0$ and $p < p' < 1$ such that $\tilde{p}_m(t) \leq C(t)(p')^m$ for any $m \geq 1$. Then we immediately obtain

$$\sum_{m=n}^{\infty} \tilde{p}_m(t) \leq C(t)\frac{(p')^n}{1-p'}.$$

Now we return to the coupled feedback loop with reaction scheme

$$\begin{gathered} G \xrightarrow{a+\mu n} G^*, \quad G^* \xrightarrow{b+\nu n} G, \\ G^* \xrightarrow{\rho p^k(1-p)} G^* + kP, \ \ k \geq 1, \quad P \xrightarrow{d} \varnothing. \end{gathered} \tag{2}$$

Note that protein molecules are produced only when the gene is active. Let $\tilde{X}_t$ and $X_t$ denote the protein number at time $t$ for the simple model given in Eq. (1) and the more complicated model given in Eq. (2) respectively. It is clear that the two Markovian models can be constructed in the same probability space so that $\tilde{X}_t \geq X_t$ for any $t \geq 0$. This implies that

$$p_n(t) \leq \mathbb{P}\left(X_t \geq n\right) \leq \mathbb{P}\left(\tilde{X}_t \geq n\right) = \sum_{m=n}^{\infty} \tilde{p}_m(t) \leq C(t)\frac{(p')^n}{1-p'}.$$

Hence, letting $K(t) = C(t)/(1-p')$ and $\gamma = -\log p'$, we have

$$p_n(t) \leq K(t)e^{-\gamma n}.$$

This indicates that $p_n(t)$ must decay exponentially with respect to the protein number $n$ when $n \geq 1$.

### 2 Proof that Eq. (15) in the main text cannot be holomorphic at both $z = z_0$ and $z = 1$

In the main text, we have proved that Eq. (10) has the following two linearly independent solutions:

$$(z-1)\, Hl\left(\xi, q_1; \alpha_1, \beta_1, \gamma, 2-\delta; \frac{\mu+\nu+d}{\mu+\nu}(z-z_0)\right), \tag{3}$$

$$(z-z_0)^{1-\gamma}\,(z-1)\, Hl\left(\xi, q_2; \alpha_2, \beta_2, 2-\gamma, 2-\delta; \frac{\mu+\nu+d}{\mu+\nu}(z-z_0)\right), \tag{4}$$

Next we will proved that the function given in Eq. (4) cannot be holomorphic at both $z = z_0$ and $z = 1$.

First, it is easy to see that the function given in Eq. (4) is holomorphic at $z = z_0$ if and only if $1-\gamma$ is a nonnegative integer. Moreover, similarly to the proof in Sec. 3.1, the function given in Eq. (4) is holomorphic at $z = 1$ if and only if $\lambda$ satisfies the following continued fraction equation:

$$q_2 = \frac{R_0^{(2)}P_1^{(2)}}{(Q_1^{(2)}+q_2)-}\frac{R_1^{(2)}P_2^{(2)}}{(Q_2^{(2)}+q_2)-}\frac{R_2^{(2)}P_3^{(2)}}{(Q_3^{(2)}+q_2)-}\cdots, \tag{5}$$

where

$$\begin{aligned} P_n^{(2)} &= (n-1+\alpha_2)(n-1+\beta_2), \quad R_n^{(2)} = \xi(n+1)(n+2-\gamma), \\ Q_n^{(2)} &= n((n+1-\gamma)(1+\xi)+\xi(2-\delta)+\alpha_2+\beta_2+\gamma+\delta-3), \end{aligned}$$

with the parameters $q_2$, $\alpha_2$, $\beta_2$, $\gamma$, and $\delta$ being all functions of $\lambda$ (see Eqs. (12) and (13) in the main text). However, it is not difficult to check that

$$\{\lambda:\ 1-\gamma \text{ is a positive integer}\} \cap \{\lambda:\ \lambda \text{ satisfies Eq. (5)}\} = \emptyset.$$

This implies that the function given in Eq. (4) cannot be holomorphic at both $z = z_0$ and $z = 1$.

### 3 Analytical time-dependent solution for unregulated genes

We focus on the time-dependent protein distributions for an unregulated gene. If a gene is unregulated, then its expression dynamics can be described by the reaction scheme

$$G \underset{b}{\overset{a}{\rightleftharpoons}} G^*, \quad G^* \xrightarrow{\rho p^k(1-p)} G^* + kP, \quad k \geq 1, \quad P \xrightarrow{d} \varnothing.$$

This is a special case of Eq. (2) when the positive and negative feedback strengths both vanish, i.e. $\mu = \nu = 0$. We emphasize that for an unregulated gene, the analytical time-dependent distributions of protein numbers have been derived in [3, 4]. Here we will provide a different explicit expression of the time-dependent solution. We first state the main results in this section and put the detailed proof in Sec. 4.

In the main text, we have shown that the generating functions $f_i$ and $f$ have the following series form:

$$f_i(t,z) = \sum_{n=0}^{\infty} C_n e^{\lambda_n t}\tilde{f}_i^{\lambda_n}(z), \quad f(t,z) = \sum_{n=0}^{\infty} C_n e^{\lambda_n t}\tilde{f}^{\lambda_n}(z). \tag{6}$$

When $\mu = \nu = 0$, the expressions of the eigenvalues $\lambda_n$, eigenfunctions $\tilde{f}^{\lambda_n}$, and coefficients $C_n$ in Eq. (6) can be simplified to a great extent. In this case, the related parameters in Eq. (18) in the main text reduce to

$$z_0 = 1, \quad w = B, \quad c_1 = \frac{a+b}{d}, \quad a_1 + b_1 = \frac{a+b+\rho}{d}, \quad a_1 b_1 = \frac{a\rho}{d^2}. \tag{7}$$

It can be proved that the system has two families of eigenvalues (see Section 4 for details)

$$\lambda_{1n} = -nd, \quad \lambda_{2n} = -(a+b+nd), \quad n \geq 0, \tag{8}$$

which can be computed explicitly. For the family of eigenvalues $\lambda_{1n} = -nd$, the corresponding eigenfunctions are given by

$$\tilde{f}^{\lambda_{1n}}(z) = (z-1)^n y_1(z),$$

and for the other family of eigenvalues $\lambda_{2n} = -(a+b+nd)$, the corresponding eigenfunctions are given by

$$\tilde{f}^{\lambda_{2n}}(z) = (z-1)^{n+1} y_2(z),$$

where

$$y_1(z) = {}_2F_1(a_1, b_1; c_1; B(z-1)), \quad y_2(z) = {}_2F_1(a_1+1-c_1, b_1+1-c_1, 2-c_1; B(z-1)),$$

and $a_1$, $b_1$, and $c_1$ are given in Eq. (15). Hence for an unregulated gene, the generating function $f$ has the form of

$$f(t,z) = \sum_{n=0}^{\infty} C_{1n} e^{-ndt} (z-1)^n y_1(z) + \sum_{n=0}^{\infty} C_{2n} e^{-(a+b+nd)t} (z-1)^{n+1} y_2(z).$$

Finally, we show how to determine the coefficients $C_{1n}$ and $C_{2n}$. Let $F_n^i$ be be the unnormalized $n$th factorial moment of the protein number at time $t = 0$ when the gene is in state $i$, i.e.

$$F_n^i = \sum_{m=0}^{\infty} m(m+1)\cdots(m+n-1) p_{i,m}(0) = \frac{\partial^n f_i}{\partial z^n}(0,z)\Big|_{z=1}. \tag{9}$$

It is easy to see that $C_{10} = 1$ and the remaining coefficients can be computed inductively as follows (see Section 4 for details):

$$\begin{aligned} C_{1n} &= \frac{F_n^0 + F_n^1 - \sum_{m=0}^{n-1} C_{1m} m! \zeta_{n-m} - \sum_{m=0}^{n-1} C_{2m}(m+1)! \xi_{n-m-1}}{n!}, \quad n \geq 1, \\ C_{2n} &= \frac{\sum_{m=0}^{n} C_{1m} m! \kappa_{n-m} - \sum_{m=0}^{n-1} C_{2m} m! \eta_{n-m} + \sum_{m=0}^{n-1} C_{2m}(m+1)! \theta_{n-m-1} - \rho p F_n^1}{(a+b-d)(1-p)n!}, \quad n \geq 0. \end{aligned} \tag{10}$$

where

$$\zeta_n = \frac{B^n (a_1)_n (b_1)_n}{(c_1)_n}, \quad \xi_n = \frac{B^n (a_1+1-c_1)_n (b_1+1-c_1)_n}{(2-c_1)_n},$$
$$\kappa_n = (1-p)\zeta_{n+1} - np\zeta_n, \quad \eta_n = (a+b-d)[(1-p)\xi_n - np\xi_{n-1}], \quad \theta_n = d[(1-p)\xi_{n+1} - np\xi_n].$$

Now we focus on the initial condition in Sec. 4 in the main text, i.e. initially the protein number is zero and the gene is inactive. In this case, we have $f_0(0,z) = 1$ and $f_1(0,z) = 0$. This clearly shows that $F_0^0 = 1$, $F_n^0 = 0$ for all $n \geq 1$, and $F_n^1 = 0$ for all $n \geq 0$. Inserting these values of $F_n^i$ into Eq. (10) gives the values of all the coefficients $C_{1n}$ and $C_{2n}$.

### 4 Proof of the results in Section 3

When the gene is unregulated ($\mu = \nu = 0$), it is easy to check that $\xi = \infty$ in Eq. (11) in the main text. Hence we cannot directly obtain the simplified expressions by taking $\mu = \nu = 0$ in Eq. (22) in the main text.

We first compute the eigenvalues and the corresponding eigenfunctions in closed form. Clearly, when the gene is unregulated, $\lambda = 0$ is still an eigenvalue. Next we determine all nonzero eigenvalues. Recall that $\lambda$ is a nonzero eigenvalue if and only if $\lambda$ satisfies the following continued fraction equation

$$q_1 = \frac{R_0^{(1)} P_1^{(1)}}{(Q_1^{(1)} + q_1) -} \frac{R_1^{(1)} P_2^{(1)}}{(Q_2^{(1)} + q_1) -} \frac{R_2^{(1)} P_3^{(1)}}{(Q_3^{(1)} + q_1) -} \cdots, \tag{11}$$

where $q_1$ and other parameters are given in Eqs. (12) and (13) in the main text, and

$$\begin{aligned} R_n^{(1)} &= \xi(n+1)(n+\gamma), \quad P_n^{(1)} = (n-1+\alpha_1)(n-1+\beta_1), \\ Q_n^{(1)} &= n[(n-1+\gamma)(1+\xi)+\xi(2-\delta)+\alpha_1+\beta_1-\gamma+\delta-1]. \end{aligned}$$

Let

$$d_n = -\frac{R_{n-1}^{(1)}P_n^{(1)}}{\xi^2}, \quad n \geq 1, \quad e_n = \frac{Q_n^{(1)}+q_1}{\xi}, \quad n \geq 0,$$

and let $C_{nm}$ be the continued fraction defined by

$$C_{nm} = e_n + \frac{d_{n+1}}{e_{n+1}+}\frac{d_{n+2}}{e_{n+2}+}\cdots\frac{d_m}{e_m} := \frac{D_{nm}}{E_{nm}}, \quad 0 \leq n < m \leq \infty,$$

where $D_{nm}$ and $E_{nm}$ are (canonical) numerator and denominator, respectively. For example, it is easy to see that

$$C_{n,n+2} = e_n + \frac{d_{n+1}}{e_{n+1}+}\frac{d_{n+2}}{e_{n+2}} = \frac{e_ne_{n+1}e_{n+2}+e_nd_{n+2}+e_{n+2}d_{n+1}}{e_{n+1}e_{n+2}+d_{n+2}}.$$

This implies that $D_{n,n+2} = e_ne_{n+1}e_{n+2}+e_nd_{n+2}+e_{n+2}d_{n+1}$ and $E_{n,n+2} = e_{n+1}e_{n+2}+d_{n+2}$. It follows from [5, Sec.1.12] that

$$D_{nm} = e_mD_{n,m-1}+d_mD_{n,m-2}, \quad E_{nm} = e_mE_{n,m-1}+d_mE_{n,m-2}, \quad m \geq n+1, \tag{12}$$

where $D_{n,n-1} = 1$, $D_{nn} = e_n$, $E_{n,n-1} = 0$, and $E_{nn} = 1$. With the above notation, the continued fraction equation given in Eq. (11) can be rewritten as $C_{0\infty} = 0$.

When $\mu, \nu \to 0$, it is clear that $\xi \to \infty$. Moreover, we have

$$\lim_{\mu,\nu\to\infty} d_n = 0, \quad \lim_{\mu,\nu\to\infty} e_n = \frac{[\lambda+(n+1)d](\lambda+a+b+nd)}{d^2}. \tag{13}$$

Combining Eqs. (12) and (13), we have

$$\lim_{\mu,\nu\to 0} D_{nm} = \lim_{\mu,\nu\to 0} e_mD_{n,m-1}, \quad \lim_{\mu,\nu\to 0} E_{nm} = \lim_{\mu,\nu\to 0} e_mE_{n,m-1}, \quad m \geq n+1.$$

If $\lambda \neq -(n+1)d$ and $\lambda \neq -a-b-nd$ for some $n \geq 0$, it is easy to see that for $m \geq n$,

$$\lim_{\mu,\nu\to 0} C_{nm} = \frac{\lim_{\mu,\nu\to 0} D_{nm}}{\lim_{\mu,\nu\to 0} E_{nm}} = \cdots = \frac{\lim_{\mu,\nu\to 0} D_{nn}}{\lim_{\mu,\nu\to 0} E_{nn}} = \lim_{\mu,\nu\to 0} e_n = \frac{[\lambda+(n+1)d](\lambda+a+b+nd)}{d^2}. \tag{14}$$

For any fix $\lambda$, let $n_\lambda = \max\{\lfloor -\mathrm{Re}(\lambda)/d\rfloor, \lfloor -\mathrm{Re}(\lambda+a+b)/d\rfloor+1\}$, where $\lfloor a\rfloor$ is the largest integer that is less than or equal to $a$. From Eq. (13), if $\mu$ and $\nu$ are sufficiently small, then

$$|e_n| \geq |d_n|+1, \quad n \geq n_\lambda+1.$$

It follows from Pringsheim's theorem [5, Sec.1.12] that when $\mu$ and $\nu$ are sufficiently small, $\lim_{m\to\infty} C_{n_\lambda m}$ exists and we set $C_{n_\lambda\infty} = \lim_{m\to\infty} C_{n_\lambda m}$. Thus it follows from Eq. (14) that

$$\lim_{\mu,\nu\to 0} C_{n_\lambda\infty} = \lim_{\mu,\nu\to 0} e_{n_\lambda} = \frac{[\lambda+(n_\lambda+1)d](\lambda+a+b+n_\lambda d)}{d^2}.$$

Furthermore, when $\mu, \nu \to 0$, we have

$$C_{0\infty} = e_0 + \frac{d_1}{e_1+}\frac{d_2}{e_2+}\cdots\frac{d_{n_\lambda}}{C_{n_\lambda\infty}} \approx e_0 + \frac{d_1}{e_1+}\frac{d_2}{e_2+}\cdots\frac{d_{n_\lambda}}{e_{n_\lambda}} = C_{0n_\lambda}.$$

If $\lambda \neq -(n+1)d$ and $\lambda \neq -a-b-nd$ for some $n \geq 0$, letting $n=0$ and $m=n_\lambda$ in Eq. (14), we have

$$\lim_{\mu,\nu\to 0} C_{0,\infty} = \lim_{\mu,\nu\to 0} C_{0n_\lambda} = \lim_{\mu,\nu\to 0} e_0 = \frac{(\lambda+d)(\lambda+a+b)}{d^2} \neq 0.$$

This implies that if $\lambda \neq -(n+1)d$ and $\lambda \neq -a-b-nd$ for some $n \geq 0$, then $\lambda$ is not an eigenvalue. On the other hand, it is easy to check that $-(n+1)d$ and $-a-b-nd$, $0 \leq n \leq N$, are all roots of the limiting equation

$$\lim_{\mu,\nu\to 0} C_{0N} = 0.$$

This implies that $-(n+1)d$ and $-a-b-nd$, $n \geq 0$, are all nonzero eigenvalues of the system. In summary, we have the following two families of eigenvalues:

$$\lambda_{1n} = -nd, \quad \lambda_{2n} = -(a+b+nd), \quad n \geq 0.$$

We next determine the eigenfunctions corresponding to all eigenvalues. For the zero eigenvalue $\lambda_{10}=0$, it follows from Eq. (20) in the main text that the eigenfunction reduces to

$$\tilde{f}^{\lambda_{10}}(z) = {}_2F_1(a_1,b_1;c_1;B(z-1)),$$

where

$$a_1+b_1 = \frac{a+b+\rho}{d}, \quad a_1b_1 = \frac{a\rho}{d^2}, \quad c_1 = \frac{a+b}{d}. \tag{15}$$

On the other hand, recall the following identity for local Heun functions [6]:

$$Hl\left(\xi,q;\alpha,\beta,\gamma,\delta;x\right) = Hl\left(\frac{1}{\xi},\frac{q}{\xi};\alpha,\beta,\gamma,\alpha+\beta-\gamma-\delta+1;\frac{x}{\xi}\right).$$

Let

$$\xi_4 = \frac{1}{\xi}, \quad q_4 = \frac{q_1}{\xi}, \quad \delta_4 = \alpha_1+\beta_1-\gamma+\delta-1, \quad \epsilon_4 = 2-\delta.$$

It then follows from Eq. (20) in the main text that for any nonzero eigenvalues $\lambda_n$, $n \geq 1$, we have

$$\tilde{f}^{\lambda_n}(z) = (z-1)Hf\left(\xi_4,q_4;\alpha_1,\beta_1,\gamma,\delta_4;\frac{\mu+\nu+d}{(\mu+\nu)\xi}(z-z_0)\right). \tag{16}$$

Let $y=(\mu+\nu+d)(z-z_0)/((\mu+\nu)\xi)$. Recall that if $|y| < \min\{|\xi_4|,1\}$, the Heun function has the Maclaurin expansion [5, Sec.31.3]

$$Hf\left(\xi_4,q_4;\alpha_1,\beta_1,\gamma,\delta_4;y\right) = \sum_{n=0}^{\infty} l_n y^n. \tag{17}$$

where

$$\xi_4\gamma l_1 - q_4 l_0 = 0, \quad R'_n l_{n+1} - (Q'_n+q_4)l_n + P'_n l_{n-1} = 0, \quad n \geq 1, \tag{18}$$

with

$$P'_n = (n-1+\alpha_1)(n-1+\beta_1), \quad Q'_n = n((n-1+\gamma)(1+\xi_4)+\xi_4\delta_4+\epsilon_4), \quad R'_n = \xi_4(n+1)(n+\gamma).$$

When $\mu,\nu \to 0$, it is easy to see that $y \to B(z-1)$ and

$$\begin{aligned} &\xi_4 \to 0, \quad q_4 \to \frac{(\lambda+d)(\lambda+a+b)}{d^2}, \quad \gamma \to \frac{\lambda+a+b}{d}, \quad \delta_4 \to \frac{d+\rho}{d}, \quad \epsilon_4 = 2+\frac{\lambda}{d}, \\ &\alpha_1\beta_1 \to \frac{a\rho+(\lambda+d)(a+b+\rho)+(\lambda+d)^2}{d^2}, \quad \alpha_1+\beta_1 \to \frac{a+b+\rho+2d+2\lambda}{d}. \end{aligned} \tag{19}$$

Hence, when $\mu = \nu = 0$, Eq. (18) reduces to $(\lambda + d)(\lambda + a + b)c_0 = 0$ and

$$\frac{[\lambda + (n+1)d](\lambda + a + b + nd)}{d^2} l_n = (n - 1 + \alpha_1)(n - 1 + \beta_1) l_{n-1}, \quad n \geq 1. \tag{20}$$

Suppose that $\lambda = -(n+1)d$ for some $n \geq 0$. It then follows from Eq. (20) that $l_0 = l_1 = \cdots = l_{n-1} = 0$ and

$$l_{n+m} = \frac{(\alpha_1 + n)_m (\beta_1 + n)_m}{m!((a+b)/d)_m} = \frac{(a_1)_m (b_1)_m}{m!(c_1)_m} l_n, \quad m \geq 0,$$

where $a_1$, $b_1$, and $c_1$ are the parameters given in Eq. (15) and $(a)_n = a(a+1)\cdots(a+n-1)$ denotes the Pochhammer symbol. Without loss of generality, we set $l_n = 1$. Recall the definition of Gaussian hypergeometric function [5, Sec.15.2]. Then we immediately obtain

$$\lim_{\mu,\nu \to 0} \sum_{m=0}^{\infty} l_m y^m = \lim_{\mu,\nu \to 0} y^n \ {}_2F_1(a_1, b_1; c_1; y) = B^n (z-1)^n \ {}_2F_1(a_1, b_1; c_1; B(z-1)). \tag{21}$$

Combining Eqs. (16), (17), and (21), we find that the eigenfunction corresponding to the eigenvalue $\lambda_{1n} = -nd$ is given by

$$\tilde{f}^{\lambda_{1n}}(z) = (z-1)^n y_1(z),$$

where $y_1(z) = {}_2F_1(a_1, b_1; c_1; B(z-1))$. Similarly, it is easy to see that the eigenfunction corresponding to the eigenvalue $\lambda_{2n} = -(a + b + nd)$ is given by

$$\tilde{f}^{\lambda_{2n}}(z) = (z-1)^{n+1} y_2(z),$$

where $y_2(z) = {}_2F_1(a_1 + 1 - c_1, b_1 + 1 - c_1; 2 - c_1; B(z-1))$. The proof is similar and thus is omitted here.

Finally, we use the initial conditions to determine the coefficients $C_{1n}$ and $C_{2n}$. In Sec. 3.1 in the main text, we have shown that the approximated values of the coefficients can be obtained by solving Eq. (24). For an unregulated gene, we can obtain the precise values of the coefficients. Note that

$$\begin{aligned} f(t,z) &= \sum_{n=0}^{\infty} \left( C_{1n} e^{\lambda_{1n} t} \tilde{f}^{\lambda_{1n}}(z) + C_{2n} e^{\lambda_{2n} t} \tilde{f}^{\lambda_{2n}}(z) \right), \\ f_i(t,z) &= \sum_{n=0}^{\infty} \left( C_{1n} e^{\lambda_{1n} t} \tilde{f}_i^{\lambda_{1n}}(z) + C_{2n} e^{\lambda_{2n} t} \tilde{f}_i^{\lambda_{2n}}(z) \right). \end{aligned} \tag{22}$$

From [5, Sec. 15.5], we define

$$\zeta_n = y_1^{(n)}(1) = \frac{B^n (a_1)_n (b_1)_n}{(c_1)_n}, \quad \xi_n = y_2^{(n)}(1) = \frac{B^n (a_1 + 1 - c_1)_n (b_1 + 1 - c_1)_n}{(2 - c_1)_n}.$$

To proceed, let $F_n^i$ be be the unnormalized $n$th factorial moment of the protein number at time $t = 0$ when the gene is in state $i$, as defined in Eq. (9). It follows from Eq. (22) that

$$F_n^0 + F_n^1 = \frac{\partial^n f}{\partial z^n}(0,z)\Big|_{z=1} = \sum_{m=0}^{n} C_{1m} m! \zeta_{n-m} + \sum_{m=0}^{n-1} C_{2m}(m+1)! \xi_{n-1-m}. \tag{23}$$

Taking $n = 0$ in Eq. (23), it is easy to see that $C_{10} = 1$. Furthermore, it follows from Eq. (23) that

$$C_{1n} = \frac{F_n - \sum_{m=0}^{n-1} C_{1m} m! \zeta_{n-m} - \sum_{m=0}^{n-1} C_{2m}(m+1)! \xi_{n-1-m}}{n!}, \quad n \geq 1.$$

This provides the iterative relation between $C_{1n}$ and the first $2(n-1)$ terms $C_{10}, C_{20}, \cdots, C_{1,n-1}, C_{2,n-1}$. To calculate the remaining coefficients, we need the iterative relation for $C_{2n}$. Similarly, it follows from Eq. (22) that

$$F_n^1 = \frac{\partial^{(n)} f_1}{\partial z^{(n)}}(0, z)|_{z=1} = \sum_{m=0}^{\infty} C_{1m} (\tilde{f}_1^{\lambda_{1m}})^{(n)}(1) + \sum_{m=0}^{\infty} C_{2m} (\tilde{f}_1^{\lambda_{2m}})^{(n)}(1).$$

Recall that

$$\tilde{f}_1^{\lambda}(z) = \frac{1 - pz}{\rho p(z-1)} \left[ \lambda \tilde{f}^{\lambda}(z) + d(z-1)\partial_z \tilde{f}^{\lambda}(z) \right].$$

It is easy to check that

$$\rho p F_n^1 = \sum_{m=0}^{n} C_{1m} m! \kappa_{n-m} - \sum_{m=0}^{n} C_{2m} m! \eta_{n-m} + \sum_{m=0}^{n-1} C_{2m}(m+1)! \theta_{n-1-m}. \tag{24}$$

where

$$\begin{aligned}
\kappa_n &= \{(1-pz)y_1'(z)\}^{(n)}|_{z=1} = (1-p)\zeta_{n+1} - np\zeta_n, \\
\eta_n &= (a+b-d)\{(1-pz)y_2(z)\}^{(n)}|_{z=1} = (a+b-d)[(1-p)\xi_n - np\xi_{n-1}], \\
\theta_n &= d\{(1-pz)y_2'(z)\}^{(n)}|_{z=1} = d[(1-p)\xi_{n+1} - np\xi_n].
\end{aligned}$$

Then we immediately obtain

$$C_{2n} = \frac{\sum_{m=0}^{n} C_{1m} m! \kappa_{n-m} - \sum_{m=0}^{n-1} C_{2m} m! \eta_{n-m} + \sum_{m=0}^{n-1} C_{2m}(m+1)! \theta_{n-1-m} - \rho p F_n^1}{(a+b-d)(1-p)n!}, \quad n \geq 0.$$

This provides the iterative relation between $C_{2n}$ and the first $2n-1$ terms $C_{10}, \cdots, C_{1n}, C_{20}, \cdots, C_{2,n-1}$.

### 5 Reduction of the Heun function to the confluent Heun function

Recall that the local Heun function $Hl(\xi, q; \alpha, \beta, \gamma, \delta; z)$ is defined as the solution of the following equation that is holomorphic at $z = 0$ and is equal to 1 there [5, Sec.31.3]:

$$h''(z) + \left( \frac{\gamma}{z} + \frac{\delta}{z-1} + \frac{\epsilon}{z-\xi} \right) h'(z) + \frac{\alpha\beta z - q}{z(z-1)(z-\xi)} h(z) = 0, \tag{25}$$

where $\alpha$ and $\beta$ are related by $\alpha + \beta + 1 = \gamma + \delta + \epsilon$. Now we consider the following limit:

$$\xi \to \infty, \quad -\frac{\epsilon}{\xi} \to \epsilon', \quad -\frac{\alpha\beta}{\xi} \to \alpha', \quad -\frac{q}{\xi} \to q'.$$

In this limit, Eq. (25) reduces to

$$h''(z) + \left( \frac{\gamma}{z} + \frac{\delta}{z-1} + \epsilon' \right) h'(z) + \frac{\alpha' z - q'}{z(z-1)} h(z) = 0. \tag{26}$$

This is exactly the standard confluent Heun differential Equation [5, Sec.31.12]. Recall that the local confluent Heun function $Hc(q', \alpha', \gamma, \delta, \epsilon'; z)$ is defined as the solution of Eq. (26) that is holomorphic at $z = 0$ and is equal to 1 there [7]. Hence in the above limit, we have

$$Hl(\xi, q; \alpha, \beta, \gamma, \delta; z) \to Hc(q', \alpha', \gamma, \delta, \epsilon'; z).$$

It follows from Eqs. (12) and (13) in the main text that when $\rho \to \infty$ and $B \to 0$, while keeping $\rho B = s$ as constant, we have the following limit:

$$\xi \to \infty, \quad -\frac{\alpha_1 + \beta_1 - \gamma + \delta - 1}{\xi} \to \epsilon_3, \quad -\frac{\alpha_1\beta_1}{\xi} \to \alpha_3, \quad -\frac{q_1}{\xi} \to q_3.$$

Then for any nonzero eigenvalue $\lambda_n$, $n \geq 1$, the corresponding eigenfunction reduces to

$$\begin{aligned}\tilde{f}^{\lambda_n}(z) &= (z-1)Hf\left(\xi, q_1; \alpha_1, \beta_1, \gamma, 2-\delta; \frac{\mu+\nu+d}{\mu+\nu}(z-z_0)\right)\\ &\to (z-1)Hc\left(q_3, \alpha_3, \gamma_3, \delta_3, \epsilon_3, \frac{\mu+\nu+d}{\mu+\nu}(z-z_0)\right).\end{aligned}$$

### 6 Origin of the incorrect eigenvalues and comparison with FSP

Here we explain how the false eigenvalues given in [8] originate from and compare them to with the true eigenvalues obtained by solving the continued fraction equation. Note that [8] only considers the special case of negative feedback loops ($\mu = b = 0$) when protein synthesis is non-bursty. However here we focus on the full model given in Eq. (2).

In fact, the method used in [8] to extract the eigenvalues is essentially the same as that applied in Sec. 3.1. However, the authors did not deal with the holomorphism of the eigenfunctions in a correct way. Recall that the two linearly independent solutions of Eq. (10) in the main text are given by Eqs. (3) and (4). To proceed, we recall the following identity about local Heun functions [6, 9]:

$$Hl(\xi, q; \alpha, \beta, \gamma, \delta; x) = (x-1)^{1-\delta} Hl(\xi, q_1; \alpha_1, \beta_1, \gamma, 2-\delta; x).$$

Hence the two linearly independently solutions of Eq. (10) in the main text can be rewritten as

$$(z-1)^{\delta} Hl\left(\xi, q; \alpha, \beta, \gamma, \delta; \frac{\mu+\nu+d}{\mu+\nu}(z-z_0)\right), \tag{27}$$

$$(z-z_0)^{1-\gamma}(z-1)Hl\left(\xi, q_2; \alpha_2, \beta_2, 2-\gamma, 2-\delta; \frac{\mu+\nu+d}{\mu+\nu}(z-z_0)\right). \tag{28}$$

In [8], the authors only consider the holomorphism of first part of the solutions given in Eqs. (27) and (28). Note that in order for the functions $(z-1)^{\delta}$ and $(z-z_0)^{1-\gamma}$ to be holomorphic on the unit disk, both $\delta$ and $1-\gamma$ must be nonnegative integers. This gives the two families of eigenvalues

$$\lambda_{1n} = -nd, \quad \lambda_{2n} = -n(\mu+\nu+d) - (a+b) - \frac{\rho p \nu}{\mu+\nu+d-dp}, \quad n \geq 0. \tag{29}$$

In this special case of negative feedback loops ($\mu = b = 0$) and non-bursty protein synthesis ($\rho \to \infty$, $B \to 0$, while keeping $\rho B = s$ as constant), the above eigenvalues reduce to

$$\lambda_{1n} = -nd, \quad \lambda_{2n} = -(n+k)(d+\nu), \quad n \geq 0,$$

where $k = a/(d+\nu) + s\nu/(d+\nu)^2$. Note that these are exactly the incorrect eigenvalues given in Eq. (29) in the main text.

In summary, the authors in [8] only used the holomorphism of parts of the solutions given in Eqs. (27) and (28) to derive the expressions for the eigenvalues. Specifically, they only consider the holomorphism of the first part (monomial part) but ignore the holomorphism of the second part (local Heun function part). Hence these incorrect eigenvalues cannot guarantee the eigenfunctions to be holomorphic on the unit disk. In particular, the true eigenvalues, which can be computed by solving the continued fraction equation given in Eq. (11), can be complex numbers, whereas the false ones given in Eqs. (29) are all real numbers. Fig. 1 compares the real parts of the true and false eigenvalues for positive and negative autoregulatory loops (top and bottom row of the figure, respectively). There are two interesting observations here. First, we find that the real parts of the true and false eigenvalues do not differ too much (Fig. 1(a),(d)) — the major differences are hence in the imaginary parts of the eigenvalues, as shown in Fig. 1(c),(f). This is curious since the method used in [8] does not take the holomorphism of the local Heun solution term into account. Second, the false eigenvalues underestimate the relaxation time (defined as the inverse of the spectral gap, i.e. $T = 1/|\mathrm{Re}(\lambda_1)|$ with $\lambda_1$ being the first nonzero eigenvalue [10]) to the steady state for positive autoregulation, and overestimate the relaxation time for negative autoregulation. This can be seen from Fig. 1(b),(e) by looking at the nonzero eigenvalue with the smallest magnitude.

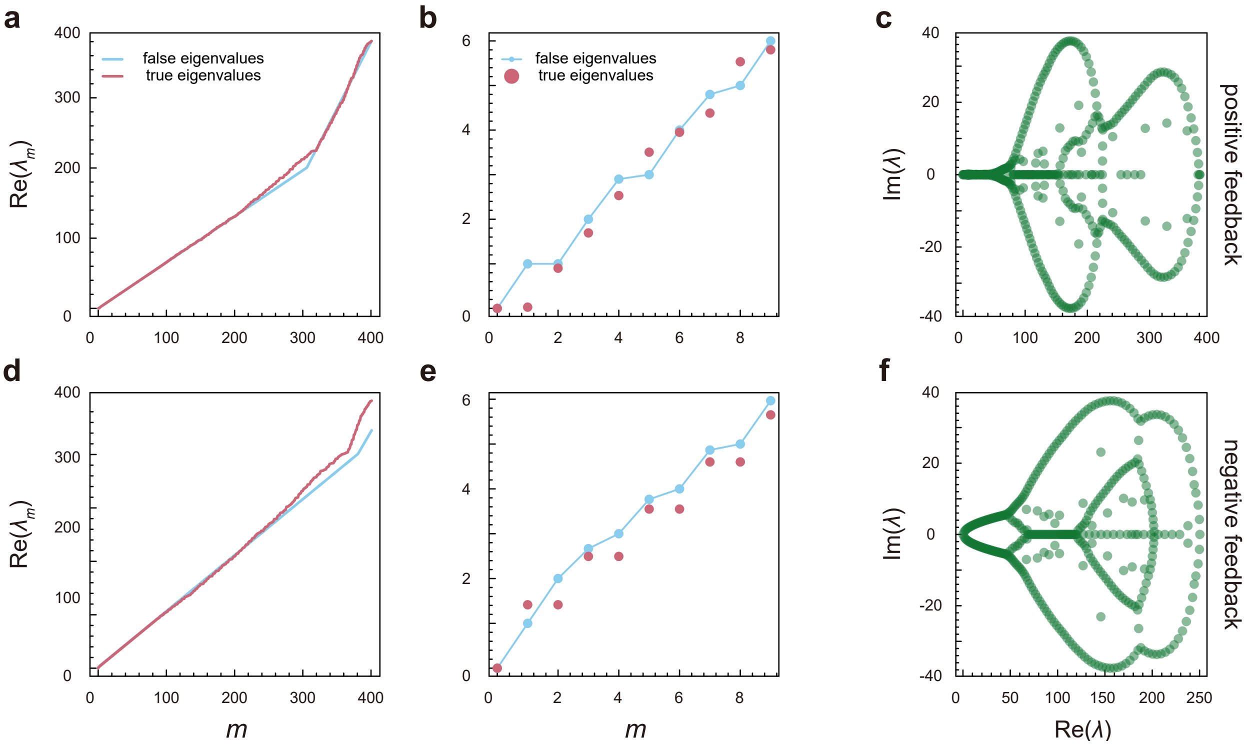


Figure 1: **Comparison of the true and false eigenvalues for positive and negative autoregulatory loops.** The true eigenvalues are computed by solving Eq. (11), and the false eigenvalues are given in Eq. (29). **(a)-(c)** Eigenvalues in positive autoregulatory loops. The parameters are chosen as $b = 1, \mu = 0.9, a = \nu = 0, \rho = 10, B = 1$. **(a),(b)** Comparison between the real parts of the true and false eigenvalues, where (a) shows the comparison for the first 400 eigenvalues and (b) shows the the comparison for the first 9 eigenvalues. **(c)** The distribution of the first 400 true eigenvalues in the complex plane. **(b)-(d)** Same as (a)-(c) but for negative autoregulatory loops. The parameters are chosen as $a = 1, \nu = 0.1, b = \mu = 0, \rho = 20, B = 1$.

### References


[1] Shahrezaei, V. & Swain, P. S. Analytical distributions for stochastic gene expression. *Proc. Natl. Acad. Sci. USA* **105**, 17256–17261 (2008).

[2] Jia, C. & Li, Y. Analytical time-dependent distributions for gene expression models with complex promoter switching mechanisms. *SIAM J. Appl. Math.* **83**, 1572–1602 (2023).

[3] Iyer-Biswas, S., Hayot, F. & Jayaprakash, C. Stochasticity of gene products from transcriptional pulsing. *Phys. Rev. E* **79**, 031911 (2009).

[4] Cao, Z. & Grima, R. Linear mapping approximation of gene regulatory networks with stochastic dynamics. *Nat. Commun.* **9**, 3305 (2018).

[5] Olver, F. W. J. *et al.* NIST Digital Library of Mathematical Functions. http://dlmf.nist.gov/, Release 1.0.17 of 2017-12-22 (2017).

[6] Maier, R. The 192 solutions of the Heun equation. *Mathematics of Computation* **76**, 811–843 (2007).

[7] Motygin, O. V. On evaluation of the confluent Heun functions. In *2018 Days on Diffraction (DD)*, 223–229 (IEEE, 2018).

[8] Ramos, A. F., Innocentini, G. C. & Hornos, J. E. M. Exact time-dependent solutions for a self-regulating gene. *Phys. Rev. E* **83**, 062902 (2011).

[9] Ronveaux, A. *Heun's differential equations* (The Clarendon Press Oxford University Press, New York, 1995).

[10] Jia, C., Qian, H., Chen, M. & Zhang, M. Q. Relaxation rates of gene expression kinetics reveal the feedback signs of autoregulatory gene networks. *J. Chem. Phys.* **148** (2018).
